# NyuWa Genome Resource: Deep Whole Genome Sequencing Based Chinese Population Variation Profile and Reference Panel

**DOI:** 10.1101/2020.11.10.376574

**Authors:** Peng Zhang, Huaxia Luo, Yanyan Li, You Wang, Jiajia Wang, Yu Zheng, Yiwei Niu, Yirong Shi, Honghong Zhou, Tingrui Song, Quan Kang, The Han100K Initiative, Tao Xu, Shunmin He

**Affiliations:** Key Laboratory of RNA Biology, Center for Big Data Research in Health, Institute of Biophysics, Chinese Academy of Sciences, Beijing 100101, China; College of Life Sciences, University of Chinese Academy of Sciences, Beijing 100049, China; National Laboratory of Biomacromolecules, CAS Center for Excellence in Biomacromolecules, Institute of Biophysics, Chinese Academy of Sciences, Beijing, 100101, China; University of Chinese Academy of Sciences, Beijing 100049, China

**Keywords:** Whole genome sequencing, Chinese population, haplotype reference panel

## Abstract

The lack of Chinese population specific haplotype reference panel and whole genome sequencing resources has greatly hindered the genetics studies in the world’s largest population. Here we presented the NyuWa genome resource based on deep (26.2X) sequencing of 2,999 Chinese individuals, and constructed NyuWa reference panel of 5,804 haplotypes and 19.3M variants, which is the first publicly available Chinese population specific reference panel with thousands of samples. Compared with other panels, NyuWa reference panel reduces the Han Chinese imputation error rate by the range of 30% to 51%. Population structure and imputation simulation tests supported the applicability of one integrated reference panel for both northern and southern Chinese. In addition, a total of 22,504 loss-of-function variants in coding and noncoding genes were identified, including 11,493 novel variants. These results highlight the value of NyuWa genome resource to facilitate genetics research in Chinese and Asian populations.

## Introduction

Comprehensive catalogues of genetic variation are fundamental building blocks in studying population and demographic history, medical genetics and genotype-phenotype association. Since the first assembly of human genome released in 2003 (International Human Genome Sequencing 2004), many large-scale whole genome sequencing (WGS) projects have been launched in Western countries and recently in Asia, and have created large and diverse population genetic variation resources. Constructing haplotype reference panel from these large cohort WGS variation resources is meaningful and cost-effective to facilitate genome-wide association study (GWAS) by imputation of unobserved genotypes into samples that have been assayed using relatively sparse genome-wide microarray chips or low coverage sequencing (Asimit and Zeggini 2012; McCarthy et al. 2016). However, as the largest ethnic group in the world, the Chinese specific reference panel is absent.

A remarkable milestone of population genome project is the 1000 Genomes Project, which released an important resource of ∼7.4X WGS data of 2,504 individuals from 26 populations, and constructed a reference panel (1KGP3) of 5,008 haplotypes at over 88 million variants (Auton et al. 2015). This resource provides a benchmark for surveys of human genetic variation, and has facilitated numerous GWASs through imputation of variants that are not directly genotyped, thus enabling a deeper understanding of the genetic architecture of complex diseases (Timpson et al. 2018). Nevertheless, rare and low-frequency variants tend to be population- or sample-specific (Auton et al. 2015), and many disease related variants are very rare and population specific (Maher et al. 2012; Saint Pierre and Genin 2014; Bomba et al. 2017). The GWASs missed a proportion of potential trait-associated variants that were poorly imputed with current reference panels (Asimit and Zeggini 2012; Hoffmann and Witte 2015; Bomba et al. 2017). So, a number of projects have focused on specific populations, attempting to capture the population specific genetic variability and build specific reference panels. For example, the Genome of the Netherlands (GoNL) Project sequenced the whole genomes of 250 Dutch parent-offspring families, found large number of novel rare variants, and constructed a reference panel with 998 haplotypes (Francioli et al. 2014). In addition, based on the GoNL panel, researchers found a rare variant rs77542162 to be associated with blood lipid levels in Dutch population (van Leeuwen et al. 2015). Afterwards there were more such projects including UK10K in United Kingdom population (Walter et al. 2015), SISu in Finnish (Chheda et al. 2017) and GenomeDenmark (Maretty et al. 2017). However, these resources are biased toward European populations. Recently some genomic resources and panels have also been created for Asian populations, including Japanese (Nagasaki et al. 2015), 219 population groups across Asia by GenomeAsia 100K project (GAsP) (Wall et al. 2019), and three Singapore populations by SG10K project (Wu et al. 2019). Some studies have also focused on Chinese population, but the sample sizes (Lan et al. 2017; Du et al. 2019) or geographical coverage (Lin et al. 2017) were limited, or genotyping methods were mainly low coverage WGS (1.7X or 0.1X) (Liu et al. 2018a; Cai et al. 2020; Gao et al. 2020). In a most recent study, the China Metabolic Analytics Project (ChinaMAP) has presented deep WGS (40.8X) dataset of 10,588 Chinese individuals mainly involved in metabolic disease (Cao et al. 2020). However, the reference panel is not yet constructed in the study. The Han Chinese population comprises about 1.23 billion people, which is the largest ethnic group in East Asia and in the world, accounting for ∼20% of the global human population and ∼ 92% of Mainland Chinese (Xu et al. 2009b). Constructing an integrated, large cohort and high quality Han Chinese population genetic variation database and reference panel is imperative, which will help to understand the population structure, population history, and facilitate genetics studies in the world’s largest population.

Here we released the genome resource named NyuWa based on high depth (median 26.2X) WGS of 2,999 Chinese individuals from 23 out of 34 administrative divisions in China. NyuWa, or Nüwa, is the mother goddess who was the creator of the human population in Chinese mythology. The NyuWa genome resource includes a total of 71.1M single nucleotide polymorphisms (SNPs) and 8.2M small insertions or deletions (indels), of which 25.0M are novel. More importantly, we constructed the NyuWa reference panel of 5,804 haplotypes and 19.3M variants, which is the first publicly available Chinese population specific reference panel with thousands of samples, and has currently the best performance for imputation of Han Chinese. We also found 1,140 pathogenic variants, 18,711 loss-of-function protein truncating variants (PTVs) and 3,793 long non-coding RNA (lncRNA) splicing variants, of which 11,493 were novel compared with existing genome resources. In a word, NyuWa genome resource can provide useful and reliable support for genetic and disease studies. The NyuWa variant database and imputation server are available at http://bigdata.ibp.ac.cn/NyuWa/.

## Results

### Large Chinese Population Cohort of Deep WGS Data

The NyuWa genome resource included high-coverage (median depth 26.2X) whole-genome sequences (WGS) of 2,999 different Chinese samples including diabetes and control samples collected from hospitals or physical examination centers. The samples were from 23 administrative divisions in China including 17 provinces, 2 autonomous regions and 4 municipalities directly under the Central Government (provinces for short, Figure 1A), which can be summarized into several geographical divisions of China (Supplementary information, Table S1). The origins of the samples were referenced to the native places or the provinces where samples were collected. The majority of samples were collected from Shanghai, Guangdong and Beijing (Figure 1A), which all have large incoming population from external provinces and rich sample diversity. The ethnicities for the samples were currently not available. Since national minorities are usually geographically clustered in China and not in our sampling areas, we estimated that the Han Chinese is the overwhelming majority in our samples.

**Figure 1.**
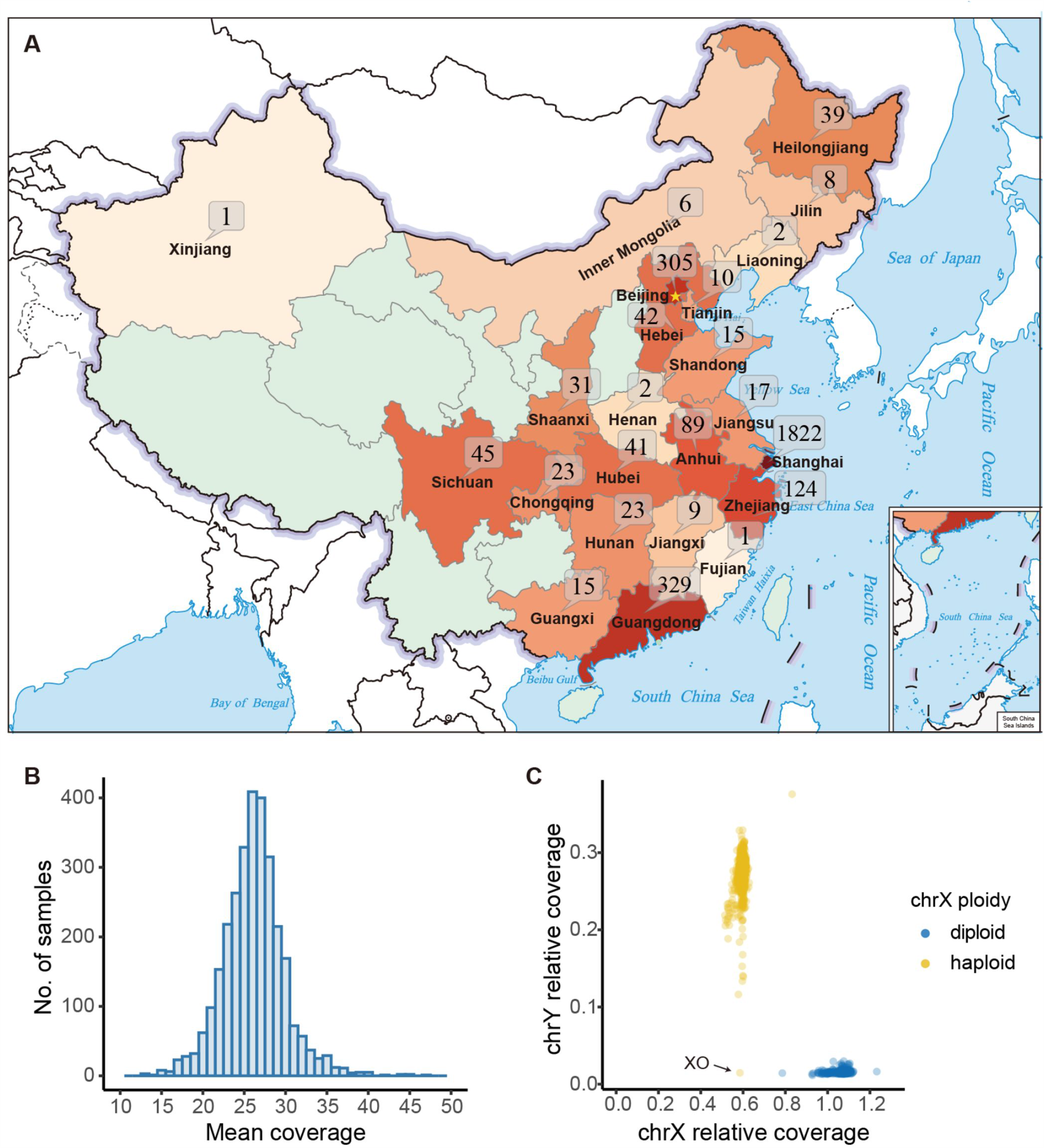
Overview of NyuWa dataset. (A) Distribution of samples in NyuWa resource. Samples were assigned to provinces based on the native places or hospitals where samples were collected. The map was downloaded from the standard map service website (http://bzdt.ch.mnr.gov.cn/). (B) The distribution of WGS mean genomic coverage after genome alignment and removal of duplicates. (C) Sex of each sample inferred by sex chromosome coverage and ploidy of chrX non-PAR region estimated by BCFtools guess-ploidy. Results were consistent for all samples except one with no chrY coverage and haploid chrX. The special sample was putative XO type and classified as female.

Most of the samples were sequenced more than 30X (median 38.9, Supplementary information, Figure S1A). After genome alignment and removal of duplicates, the median of actual genomic coverage is 26.2X (Figure 1B; Supplementary information, Figure S1B). Samples with contamination levels alpha >= 0.05 were removed (Supplementary information, Figure S1C). Based on the genomic coverage of sex chromosomes, sample sex could be clearly identified except one potential XO type (Figure 1C). The ploidy of chrX for the sample also supported the XO type, which was classified as female. There were in total 1,335 females and 1,664 males in The NyuWa dataset. After identification of close relatives within 3^rd^ degree (Supplementary information, Figure S1D), a maximum of 2,902 independent samples can be obtained in NyuWa dataset.

### 25.0M Novel Variants were Discovered in NyuWa Resource

SNPs and indels were called and genotyped using GATK cohort pipeline (Ryan Poplin 2017) with human reference genome version GRCh38/hg38 as reference. After site quality filtering, a total of 76.4M variant sites were identified, including 2.5M multi-allele sites (Supplementary information, Figure S2A). After splitting of multi-allele sites, the final dataset contained 71.1M SNPs and 8.2M indels (Supplementary information, Figure S2B), including 2.5M SNPs and 0.3M indels from sex chromosomes (Supplementary information, Table S2). The transition-to-transversion ration (Ts/Tv) is 2.107 for all bi-allelic SNPs, which is consistent with previous whole genome studies such as 1KGP3 (2.09) (Auton et al. 2015) and UK10K (2.15) (Walter et al. 2015).

Compared to other public variant repositories including ExAC (Lek et al. 2016), gnomAD (v2 & v3) (Lek et al. 2016), 1KGP3 (Auton et al. 2015), ESP (NHLBI GO Exome Sequencing Project), dbSNP (v150) (Sherry et al. 2001), GAsP (Wall et al. 2019), 90 Han (Lan et al. 2017) and TOPMed (Taliun et al. 2019), the NyuWa dataset discovered 25.0M novel variants, including 23.1M SNPs (32.5%) and 1.9M indels (23.3%) (Figure 2A). The ChinaMAP resource (Cao et al. 2020) only provided website for variant search, but did not make full variant list available. To estimate the ratio of novel variants compared with ChinaMAP, we used two variant sets for manual comparison. The first set was 230 novel singletons randomly selected from NyuWa dataset (10 per chromosome), and there were only 21.3% variants that also exist in ChinaMAP dataset. Another set was novel variants in 906 cancer related genes collected from ClinGen database and literature (Rehm et al. 2015; Huang et al. 2018; Mirabello et al. 2020). There were a total of 959k novel variants in these genes, and only 27.3% of these variants overlapping with ChinaMAP. We estimated that there were about 73% novel variants remain (∼18M) compared with ChinaMAP. As expected, most novel variants were extremely rare, with singletons, doubletons and tripletons accounting for 86.8%, 10.1%, 1.9% of novel variants, respectively (Figure 2A). This is not surprising since rare variants are usually sample- and population-specific (Francioli et al. 2014). The absolute number of novel variants with minor allele frequency (MAF) > 0.1% is still large (77.2k). These variants are frequent enough to be subject to large scale genetic association studies, and may bring new biological discoveries (Piton et al. 2013; Walter et al. 2015). The overall large number of novel variants indicates severe underrepresentation of variants in Chinese population in current genetic studies.

**Figure 2.**
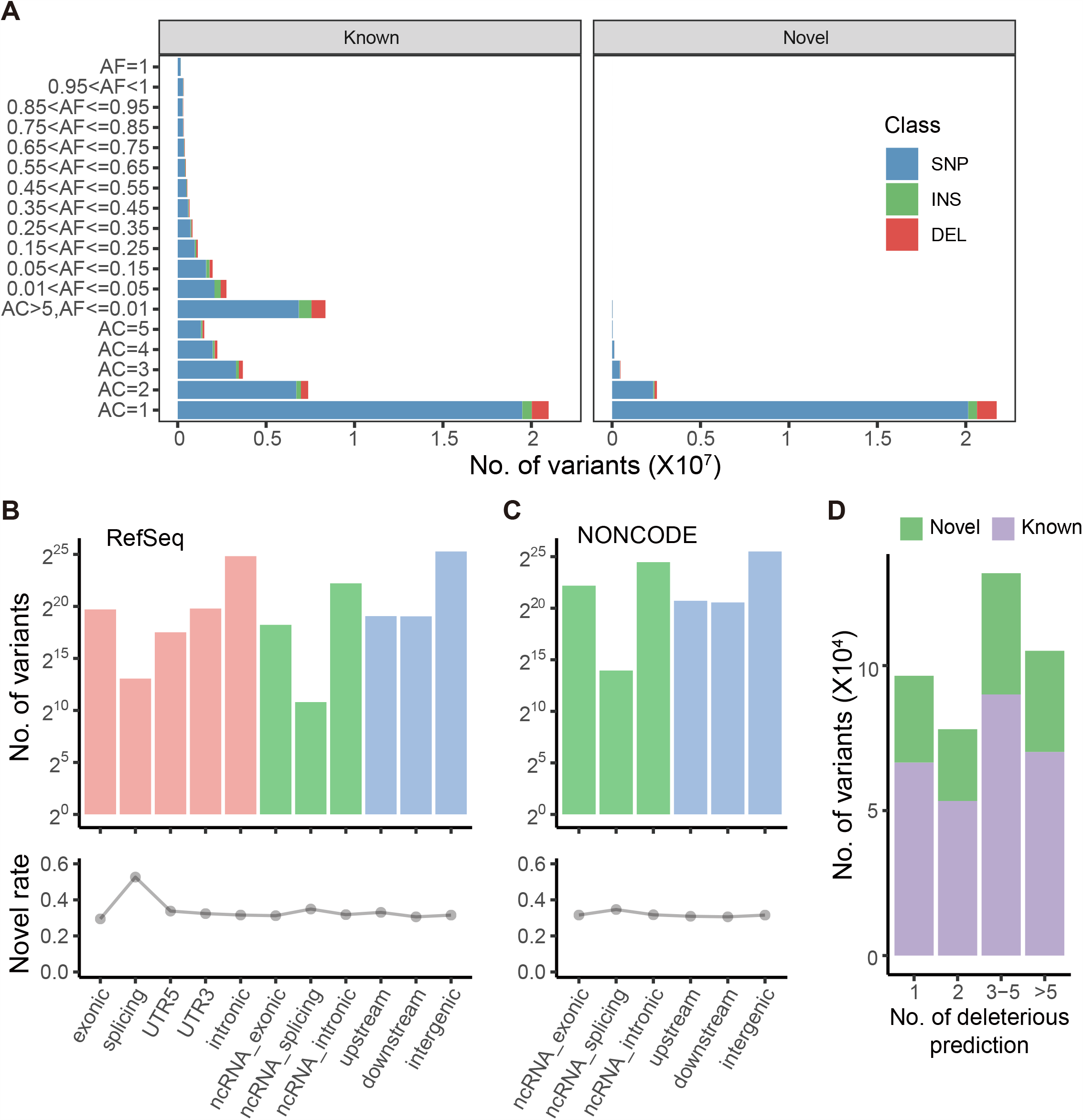
Variants statistics in NyuWa resource. (A) Number of variants detected in different bins of allele counts or frequencies. Variants were classified as known or novel based on public resources including ExAC, gnomAD v2 & v3, 1KGP3, ESP, dbSNP, TOPMed, 90 Han and GAsP. INS: small insertion, DEL: small deletion. (B) Number (upper) and novel rate (lower) of variants in different RefSeq annotation regions. (C) Number (upper) and novel rate (lower) of variants in different NONCODE annotation regions. (D) Number of non-synonymous SNPs predicted as deleterious by different number of 10 selected prediction algorithms (SIFT, Polyphen2 HDIV & HVAR, LRT, MutationTaster, FATHMM, PROVEAN, MetaSVM, MetaLR and M-CAP) provided by dbNSFP. The novel variants are based on results in (A).

On average, a NyuWa sample carries a median number of 3.51M SNPs and 523k indels in autosomes. These numbers are close to East Asia samples in 1KGP3 (3.55M SNPs, 546k indels) (Auton et al. 2015). The detected SNPs and indels with MAF > 0.1% per sample had slightly positive correlation with genomic coverage (R^2^ = 0.075 and 0.11, respectively) (Supplementary information, Figure S2C and S2D), indicating that the WGS quality can still be improved by increasing sequencing depth to higher than 30X, especially for indels. This could be explained by that although there is sufficient coverage for the whole genome, there are still regions lack coverage randomly or are difficult to amplify, which will be improved by increasing the sequencing depth. The median of MAF < 0.1% SNPs and indels in a genome were 26.4k (0.75%) and 2.57k (0.49%), respectively. The very rare SNPs and indels showed no positive correlation with sequencing depth (Supplementary information, Figure S2E and S2F), probably because the number of rare variants in different samples vary more largely (∼±10 %) compared with MAF > 0.1% variants (∼±1%), and the positive correlation is submerged by the large fluctuation.

To evaluate variant discovery by increasing sample size, we randomly down-sampled NyuWa dataset to different sizes with 100 samples intervals, and estimated the total variants and variant increase at different sample sizes (Supplementary information, Figure S2G-J). We found that the number of both SNPs and indels continued to increase with the increasing sample size (Supplementary information, Figure S2G and S2H), but the growth rate decreased, from the initial average increase of 39.4k and 5.7k per sample to the final 13.0k and 1.0k for SNPs and indels, respectively (Supplementary information, Figure S2I and S2J).

There were a total of 31.9M variants in protein coding genes, including 857k CDS, 1.10M UTR, 8.60k splicing and 30.0M intron variants (Figure 2B; Supplementary information, Figure S2K and Table S3). For lncRNAs, variants were also annotated with NONCODE v5 (Fang et al. 2017), which has the largest collection of lncRNAs. There were in total 4.78M variants in lncRNA exon regions (Figure 2C; Supplementary information, Table S4). Focusing on variants in protein coding exons, among 501k non-synonymous SNPs, 315k were annotated as deleterious by at least two of ten selected prediction algorithms provided by dbNSFP (Liu et al. 2016) (Figure 2D). The number of novel non-synonymous and deleterious SNPs were 149k and 101k, respectively (Table 1). Other functional protein coding variants included 311k synonymous SNPs, 15.3k frameshift indels, 12.7k non-frameshift indels, 11.9k stop gains and 613 stop losses (Supplementary information, Table S5). Compared to adjacent frameshift indels, there are more in-frame indels in coding region (Supplementary information, Figure S2L), consistent with previous report (Lek et al. 2016).

**Table 1.** Number of total and new variants in NyuWa resource and reference panel.

| Type | All variants <sup>a</sup> |  | Reference panel <sup>b</sup> |  |
| --- | --- | --- | --- | --- |
|  | Total | Novel <sup>c</sup> | Total | Specific <sup>d</sup> |
| All | 79,226,351 | 25,014,646 | 19,256,267 | 3,246,071 |
| Non-synonymous | 500,966 | 149,343 | 73,260 | 7,048 |
| Non-synonymous deleterious | 315,016 | 101,407 | 33,526 | 3,323 |
| **PTV** | 18,711 | 9,994 | 1,381 | 334 |
| **lncRNA splicing** | 3,793 | 1,499 | 743 | 80 |

a. All variants refers to variants in NyuWa resource. b. Reference panel refers to variants in NyuWa reference panel. c. Novelty of variants was compared with dbSNP, 1KGP3, gnomADV2.1, EXAC, ESP, gnomADV3, TOPMed, 90 Han and GAsP d. Variants in the other 4 public available haplotype reference panels (1KGP3, HRC.r1.1, GAsP and TOPMed) were excluded.

We have designed a companion database (http://bigdata.ibp.ac.cn/NyuWa_variants/) to archive SNPs and indels in NyuWa resource, and to comprehensively catalogue the variants on allele frequencies in our Chinese dataset and external datasets including 1KGP3 and gnomAD v3. Besides, variant quality metrics, genome region annotations, non-synonymous impact prediction, loss-of-function prediction, clinical annotation and pharmacogenomics annotation are also collected and presented.

### NyuWa Reference Panel Outperformed Other Publicly Available Panels for Chinese Populations

Genome-wide genotype imputation is a statistical technique to infer missing genotypes from known haplotype information, which is cost-effective for GWAS with SNP arrays when compared with whole exome sequencing (WES) or WGS. NyuWa haplotype reference panel (http://bigdata.ibp.ac.cn/refpanel/) was constructed using 19.3M SNPs and indels with minor allele count >= 5 (MAC5, approximately MAF > 0.1%) in 2,902 independent samples, including 73.3k non-synonymous and 33.5k deleterious SNPs (Table 1). Compared with 4 other publicly available reference panels including 1KGP3 (Auton et al. 2015), Haplotype Reference Consortium release 1.1 (HRC.r1.1) (McCarthy et al. 2016), GAsP (Wall et al. 2019) and TOPMed r2 (Taliun et al. 2019), NyuWa reference panel had 3.25M specific variants not included in other panels, including 7.05k non-synonymous and 3.32k deleterious SNPs (Table 1). These NyuWa panel specific variants may bring new discoveries in future association studies. To evaluate the imputation performance, array genotyping data for 58 worldwide populations from the Human Genome Diversity Project (HGDP) (Li et al. 2008) were used as a testing dataset. We focused on 16 Chinese populations and 11 other Asian populations in HGDP. NyuWa outperformed 1KGP3, HRC.r1.1 and TOPMed r2 in all Chinese populations except Uygur (Figure 3A; Supplementary information, Figure S3A and S3B). This can be explained by that the Uygur population belongs to the Central Asia and was seldom included in our sampled areas. For Han Chinese, imputation with NyuWa reduced the error rate by 38.1%, 50.8% and 30.4% compared with 1KGP3, HRC.r1.1 and TOPMed r2, respectively. NyuWa also achieved better performance in most other East Asian and Northeast Asian populations (Figure 3A; Supplementary information, Figure S3A-D). Not surprisingly, NyuWa did not perform as well as 1KGP3 in Central/South Asian populations in HGDP, which are mainly from Pakistan and historically received substantial gene flow from Central Asia and western Eurasia (Qamar et al. 2002; Majumder 2010). Comparing to GAsP, a newly released reference panel for Asian populations, NyuWa also has advantage in Chinese populations including Han, She, Tujia, Miaozu, Yizu, Tu and Naxi (Figure 3B; Supplementary information, Figure S3C). For Han Chinese, imputation with NyuWa reduced the error rate by 33.2% compared with GAsP. Nevertheless, NyuWa performed worse in some of Chinese minorities and the Pakistan Central/South Asian populations, possibly due to the overwhelming proportion of Han population in NyuWa. These results indicated that additional minority samples were needed to improve the imputation performance of certain Chinese minorities. We further compared the number of high-quality imputed variants in total imputed variants among these panels. NyuWa showed the largest number and proportion of high-quality imputed variants (Rsq > 0.8) across all MAF bins in Chinese and Han Chinese population compared with GAsP and HRC.r1.1 (Figure 3C and 3D). Compared to 1KGP3, NyuWa had larger number of high-quality imputed variants in low MAF regions (Rsq > 0.8, MAF < 0.05), and similar number in high MAF regions (MAF > 0.05) (Figure 3C and 3D). While TOPMed r2 had slightly less numbers in high MAF regions and the largest numbers in low MAF regions, the percentage of high-quality imputed variants were very low in all MAF regions, due to the largest number of total variants (an order of magnitude higher than other 4 panels) (Figure 3C and 3D).

**Figure 3.**
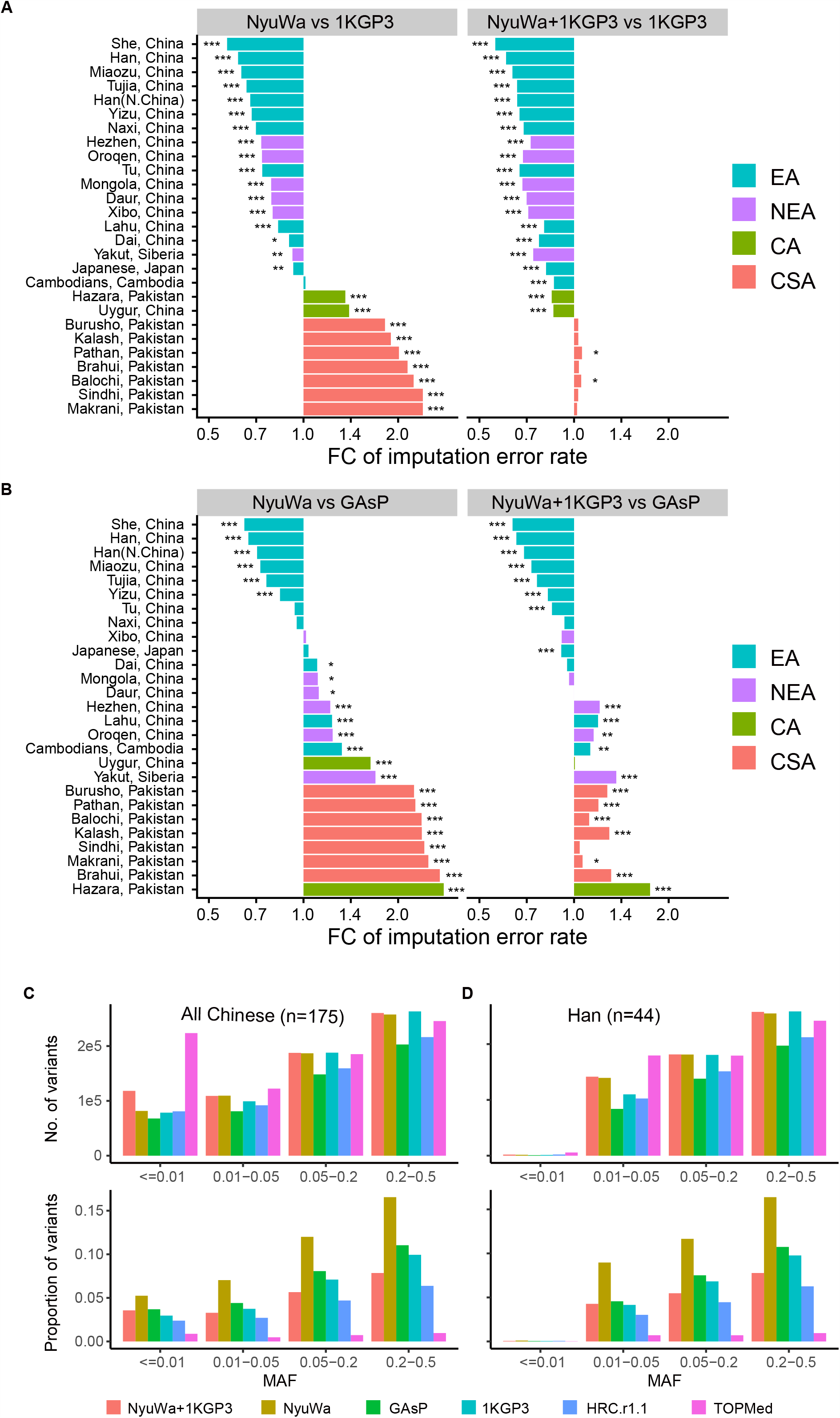
Performance of NyuWa haplotype reference panel. (A) Fold change (FC) of imputation error rate in different Asia populations in HGDP dataset between 1KGP3 panel and NyuWa (left) or NyuWa+1KGP3 (right) panel. Lower fold change values represent better performance in NyuWa or NyuWa+1KGP3 panels. EA: East Asian; NEA: North East Asian; CA: Central Asian; CSA: Central South Asian. Significances of error rate differences were calculated by chi-squared test. *: p < 0.05; **: p < 0.01; ***: p < 0.001. (B) Fold change of imputation error rate between GAsP panel and NyuWa (left) or NyuWa+1KGP3 (right) panel. Colors representing regions in (A) and (B) are consistent. (C-D) Number (upper) and proportion (lower) of high-quality (Rsq > 0.8) variants imputed for All Chinese populations (C) and Han Chinese population (D) in HGDP dataset. Variants were grouped in different MAF intervals.

To optimize imputation performance, we also combined NyuWa reference panel with 1KGP3 panel using the reciprocal imputation strategy (Huang et al. 2015). The combined panel (NyuWa + 1KGP3) included 5,406 samples and 40.2M variants, which improved imputation in all tested Asian populations (Figure 3A; Supplementary information, Figure S3). The imputation accuracy was obviously improved by about 10% for Chinese minorities of Mongolian, Dai, Daur, Xibo, Tu, Oroqen and Uygur, and outperformed GAsP in more Chinese minority populations (Figure 3B). For Chinese and Chinese Han population, NyuWa+1KGP3 could impute more high-quality variants (Rsq > 0.8) across all MAF bins, with significant increase in low MAF variants (MAF < 0.01) (Figure 3C and 3D). In brief, NyuWa+1KGP3 is an excellent alternative of NyuWa.

### Applicability of One Integrated Reference Panel for Both Northern and Southern Chinese

Due to the awareness of north-south genetic differences in Han Chinese people (Xu et al. 2009a; Chiang et al. 2018), we asked if it is adequate to use one integrated reference panel for both north and south Han populations. To answer this question, we analyzed NyuWa dataset from the perspective of population structure and imputation simulation tests.

In order to verify the ethnic authenticity of NyuWa samples, principal component analysis (PCA) of 200 randomly selected NyuWa samples together with 1KGP3 samples showed that NyuWa samples were clustered together with 1KGP3 Han Chinese samples (Supplementary information, Figure S4A and S4B), indicating that NyuWa samples are truly Chinese samples and little batch effect is observed. Y chromosome analysis of male samples in NyuWa population showed that majority (77.5%) of Y-chromosome haplogroups was O group, which is the dominant group in Han Chinese population. The following groups were C (9.0%) and N (7.5%). The Y haplogroup distribution was consistent with previous analysis of Chinese populations (Yan et al. 2014) (Supplementary information, Figure S5A). The distribution of Y haplogroups in different provinces were shown in Supplementary information, Figure S5B.

We then analyzed ancestral components of NyuWa samples. Cross validation of ADMIXTURE analysis for NyuWa with 1KGP3 East Asia samples showed that K = 3 best matched the structure of East Asia populations (Figure 4A; Supplementary information, Figure S6). Consistent with CHB (Han Chinese in Beijing, China) and CHS (Southern Han Chinese) samples in 1KGP3, the most predominant component in NyuWa samples was the ancestral component 1 (red). In the view of sample origins, a clear difference between people in north and south provinces was that south people have more proportion of ancestral component 3 (blue, Figure 4B), which was also the case between CHB and CHS samples in 1KGP3. The component 3 was also the major ancestral component for Dai (CDX) and Vietnamese (KHV) people (Figure 4A and 4B). The component 2 (green) was the major ancestral component for Japanese (JPT) people, and was minor in Chinese samples (Figure 4A and 4B).

**Figure 4.**
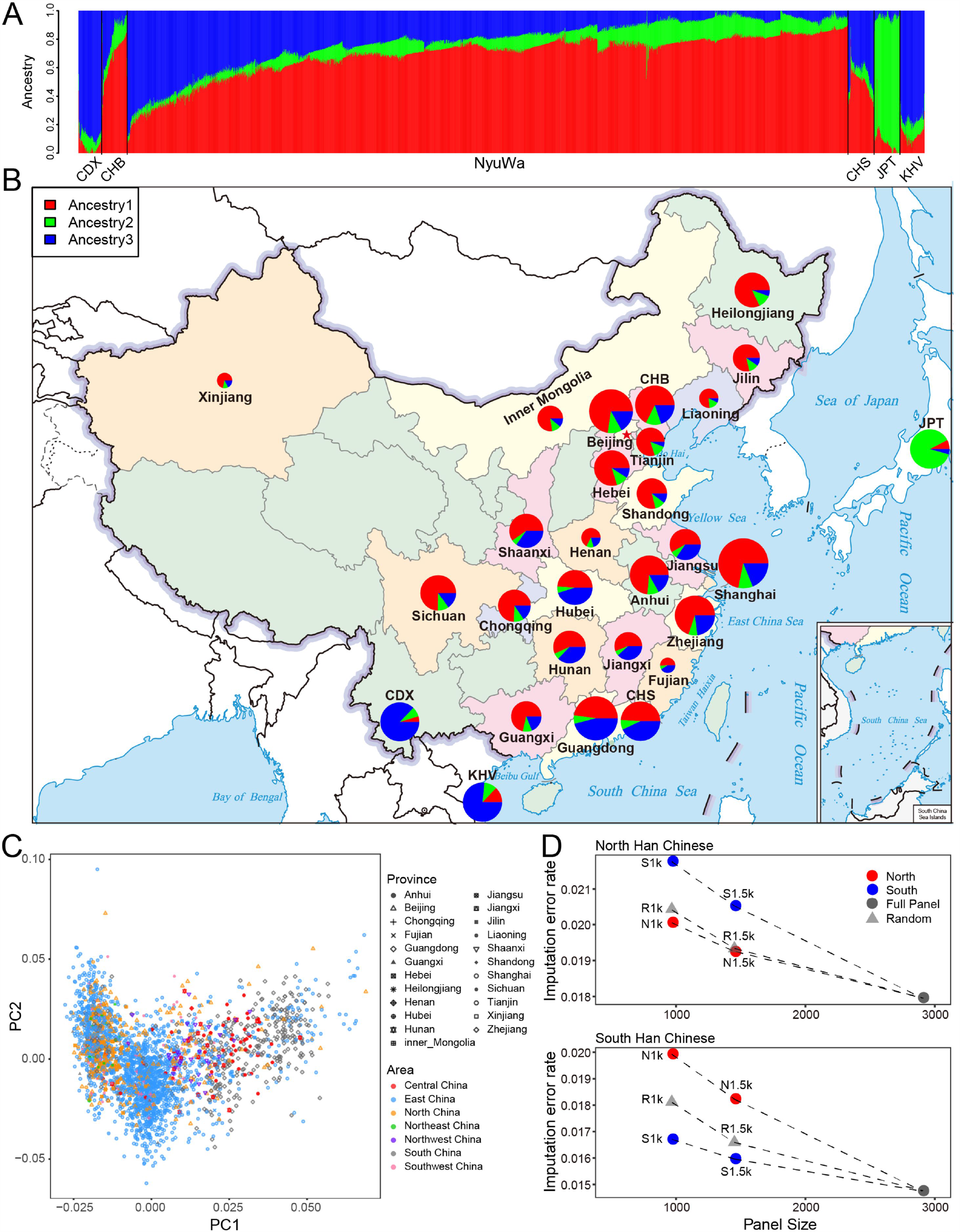
Chinese population structure based on NyuWa dataset. (A) ADMIXTURE analysis of NyuWa samples with East Asia samples in 1KGP3. Number of ancestries K = 3 best fits the model. Different colors represent different ancestry components. CHB: Han Chinese in Beijing, China; CHS: Southern Han Chinese; CDX: Chinese Dai in Xishuangbanna, China; JPT: Japanese in Tokyo, Japan; KHV: Kinh in Ho Chi Minh City, Vietnam. (B) Proportion of ancestry components in different provinces. The ancestry components and colors are consistent with (A). 1KGP3 East Asia populations (CHB, CHS, CDX, JPT and KHV) were also shown. (C) Top 2 primary components (PC1 & 2) of NyuWa samples. Each point represents a sample. Samples were marked by provinces and areas of China. PC1 represents the difference of north and south Chinese. (D) Imputation error rates of two test datasets representing north (Han N. China in HGDP, upper) and south (CHS in 1KGP3, lower) Han Chinese. Each point represents a reference panel constructed with a certain sample subset of NyuWa reference panel. The color of red represents North (N) specific panels from samples in the left part of PC1 shown in (C), while blue represent South (S) specific panels in the right part of PC1. The gray triangles represent reference panels with randomly (R) selected samples. The numbers 1k and 1.5k represent the proportions 1/3 and 1/2 of the 2902 total samples in NyuWa panel. Dotted lines represent addition of more samples.

The above ADMIXTURE results indicated that north and south Chinese share two major ancestral components, and are different at the proportions of these components, which is consistent with the historical migration and partial mix within the past two to three millennia (Wen et al. 2004; Chen et al. 2009). Using primary component analysis (PCA), we found that the primary component 1 (PC1) of NyuWa samples represented the trend of north to south differentiation (Figure 4C), which is consistent with previous studies for Han and Chinese minorities (Chiang et al. 2018; Liu et al. 2018a; Cao et al. 2020). Other PCs does not show differentiation between north and south (Supplementary information, Figure S7A). We observed that variants with high absolute weights in PC1 also showed high AF differences between ancestral components 1 and 3 (Supplementary information, Figure S7B). F_st_, another analysis for genetic differentiation between north and south NyuWa samples classified with the classic geographical demarcation of Qinling Mountains-Huaihe River, also showed that north-to-south differential sites are also different between ancestral components 1 and 3 (Supplementary information, Figure S7C). These results showed that the genetic differences already existed between the ancestries 1 and 3, which is consistent with the partial mix of ancestry components. Collectively, since north and south Chinese share the same major ancestral components, we reason that one integrated reference panel is applicable for both north and south Han Chinese.

To test the speculation, we divided samples from NyuWa reference panel into different north or south subsets based on sample positions on PC1, which represents differentiation between north and south (Figure 4C). North/south Han Chinese specific panels were then constructed using these sample subsets, and imputation error rates were compared on independent public datasets including north Han Chinese (Han North China in HGDP) and south Han Chinese (CHS, Chinese Han South in 1KGP3). As expected, given the same sample sizes, the north or south matched specific panels had lower imputation error rates than unmatched panels (Figure 4D). Panels with randomly selected samples had intermediate error rates. Increasing panel sizes always reduced error rates, no matter added samples are matched or unmatched (Figure 4D; Supplementary information, Figure S8A). The error rates of the integrated panel were always the lowest. The imputation results for Han Chinese Beijing (CHB) samples in 1KGP3 also showed lower error rates for panels with larger sizes (Supplementary information, Figure S8B), while the differences between north and south panels were not obvious, probably because there are also many south samples in CHB (Supplementary information, Figure S4B). Another classification way using geographical demarcation of Qinling Mountains-Huaihe River showed similar results (Supplementary information, Figure S8C&D). These results confirmed the applicability of one integrated panel for both northern and southern Chinese.

We also explored whether there is a difference in the introgression level of Denisovan and Neanderthal ancestries between north and south NyuWa populations (Supplementary information, Figure S9). No obvious north-south difference was found, suggesting that the introgression of Denisovan and Neanderthal ancestries occurred before the split time of north and south ancestral populations, which is far before the current population mix. Also we found no samples with high Denisovan ancestry (>3%) like that in Melanesians and Aeta (Wall et al. 2019). The top 10 highest Denisovan ancestry samples were from Shanghai (5), Beijing (2), Guangdong (1), Shaanxi (1) and Xinjiang (1), ranging from 0.42-0.45%.

### Clinical Annotations for Variants

To demonstrate the value of NyuWa resource in improving human health, we further evaluated the utility of NyuWa in disease genetic studies and medical applications. We annotated all the variants with ClinVar (Landrum et al. 2018), and found 1,140 pathogenic variants (Supplementary information, Figure S10A and S10B). As expected, most of the pathogenic variants were singletons or rare variants in NyuWa and public datasets (Figure 5A). Each sample had a median of 4 homozygous pathogenic variants and 7 heterozygous pathogenic variants (Supplementary information, Figure S10C). We noticed that there were 32 pathogenic variants with allele frequency (AF) > 1% (Figure 5A and Supplementary information, Data S1). Pathogenic variants are usually rare, especially for rare diseases, and pathogenic variants with high AFs may relate to common diseases, or their pathogenicity are subject to further examinations. We also found some variants annotated as conflicting interpretations of pathogenicity by ClinVar showing specific higher AFs in NyuWa resource (Figure 5B and Supplementary information, Data S1). For example, taking AF 1% as threshold, two variants rs182677317 and rs369849556 were annotated as conflicting for a rare disease ciliary dyskinesia, while the high AFs (> 1%) in NyuWa dataset suggested these variants may not be pathogenic (Figure 5C). These results showed that variant AFs in NyuWa dataset can provide additional reference to assist study of disease related variants.

**Figure 5.**
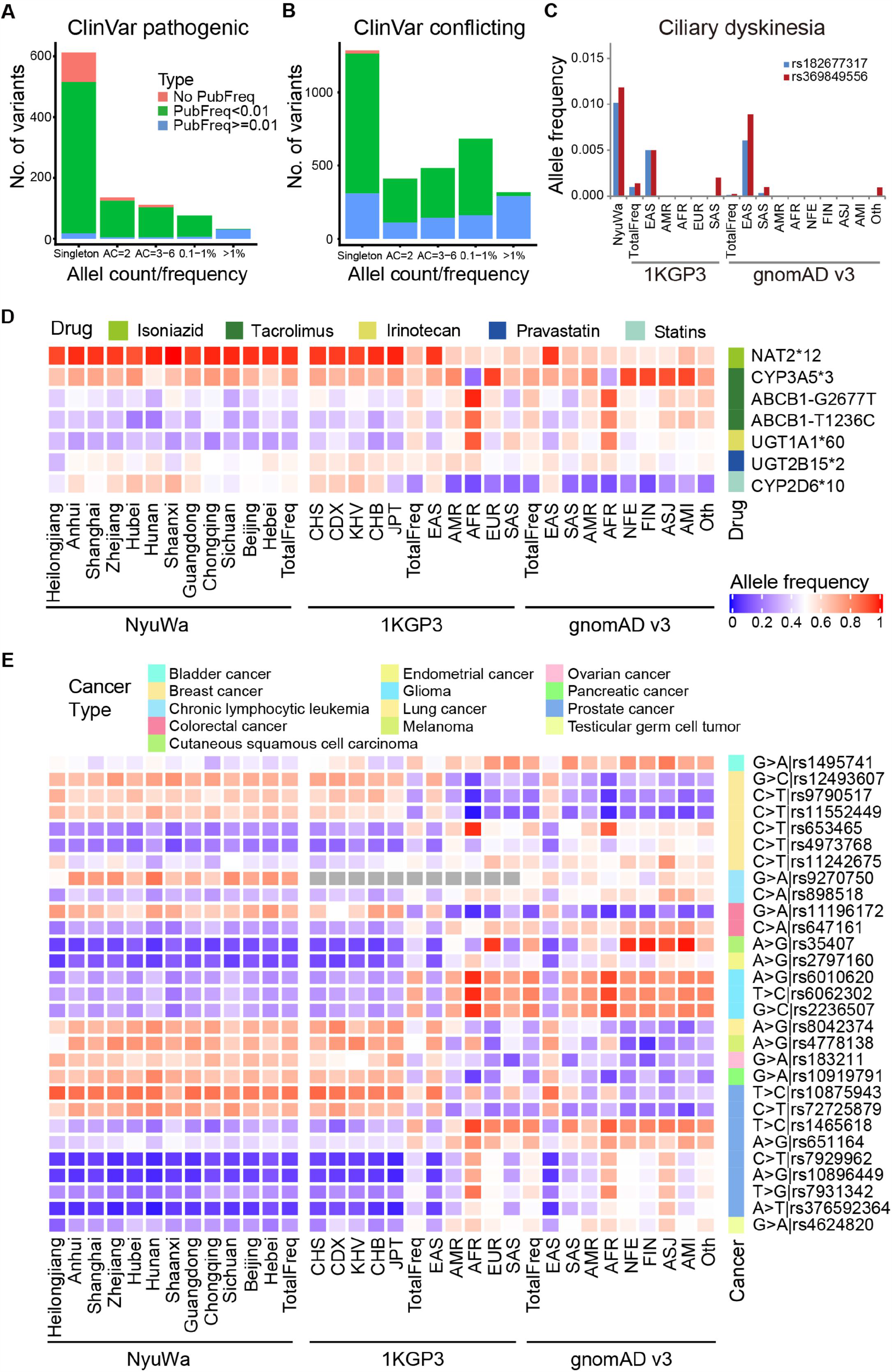
Annotation of variants. (A) Allele count and frequency distribution for ClinVar pathogenic variants. (B) Allele count and frequency distribution for ClinVar variants annotated as conflicting interpretations of pathogenicity. (C) Allele frequencies of two variants in different repositories. The two variants were annotated by ClinVar as conflicting interpretations of pathogenicity for ciliary dyskinesia. TotalFreq: the AF of all samples in the corresponding dataset; EAS: East Asian; AMR: American; AFR: African; EUR: European; SAS: South Asian; NFE: Non-Finnish European; FIN: Finnish; ASJ: Ashkenazi Jewish; AMI: Amish; Oth: Other. (D) Allele frequencies of known pharmacogenomic loci (row) that vary in different populations or regions (column). For NyuWa dataset, only provinces with sample sizes >= 20 were shown. CHB: Han Chinese in Beijing, China; CHS: Southern Han Chinese; CDX: Chinese Dai in Xishuangbanna, China; JPT: Japanese in Tokyo, Japan; KHV: Kinh in Ho Chi Minh City, Vietnam. (E) Allele frequencies of known cancer risk loci (row) that vary in different populations or regions (column). For NyuWa dataset, only provinces with sample sizes >= 20 were shown. The AF color bar is consistent with (D).

We also assessed the allele frequencies of known pharmacogenomic loci from ADME core genes (http://pharmaadme.org/) that may affect the efficacy and safety of drugs in different China provinces and worldwide regions (Supplementary information, Data S2). We found some variants with obvious AF differences in different regions of China, as well as in different populations worldwide (Figure 5D). For instance, isoniazid, a drug recommended by World Health Organization (WHO) in the treatment of tuberculosis (TB), is metabolized primarily by the NAT2 (N-Acetyltransferase 2) enzyme. *NAT2*12* refers to rs1208, and the reference allele A dampens the enzyme activity (Vatsis et al. 1991). The homozygous reference genotype will cause drug accumulation and poisoning, while heterozygous and homozygous alternative genotypes have less toxic side effects (Toure et al. 2016). We detected consistently high AFs (near 100%) of *NAT2*12* in different China provinces and East Asians, while relatively lower frequencies in other populations (Figure 5D). This suggested that testing the *NAT2*12* genotype before using isoniazid for Chinese is not as necessary as for other populations. For other examples, the AFs were not close to 0% or 100%, and vary in different China provinces (Figure 5D), hence it is recommended to take genetic tests before using certain drugs for individualized treatment.

We also checked cancer risk loci (Sud et al. 2017) in different regions (Data S2). It is generally known that there are racial differences in cancer susceptibility and survival, and the genetic factors are very important (Ozdemir and Dotto 2017). We also detected obvious AF differences between Chinese and other populations in many cancer susceptibility loci (Figure 5E).

### Loss-of-Function Variants of Protein-coding Genes and LncRNA Genes

Human loss-of-function variants have profound effects on gene function, and are informative for clinical genome interpretation. We first screened high confidence loss-of-function protein-truncating variants (PTVs), especially novel variants. We found 18,711 PTVs in 7,696 genes, in which most PTVs were singletons (Figure 6A and 6B), in line with PTV data from ExAC (67% singletons) (Lek et al. 2016). There were 9,994 novel PTVs found in NyuWa dataset, and 1,381 PTVs can be imputed by NyuWa reference panel (Table 1). The number of homozygous PTVs were 21 (Figure 6B, Supplementary information, Figure S10D). For each sample, there was a median of 24 homozygous PTVs and 58 heterozygous PTVs (Supplementary information, Figure S10E). We detected 1,138 PTVs in 385 of 906 cancer related genes, in which 636 are novel. Focusing on 9 well studied cancer-associated genes (*BRCA1, BRCA2, TP53, MEN1, MLH1, MSH2, MSH6, PMS1* and *PMS2A*) (Wall et al. 2019), there were 5 novel PTVs and 48 known ones in *BRCA2, BRCA1, PMS1, TP53* and *MSH6* (Figure 6C). Both *BRCA1* and *BRCA2* are involved in maintenance of genome stability, specifically the homologous recombination pathway for DNA double-strand break repair. Inherited mutations in *BRCA1* and *BRCA2* confer increased lifetime risk of developing breast or ovarian cancer. There were 10 known PTVs in *BRCA1* and *BRCA2*, in which 9 have been annotated as pathogenic and related to breast-ovarian cancer in ClinVar (Landrum et al. 2018), and 1 has not been collected in dbSNP yet. The uncollected and novel PTVs in *BRCA1* and *BRCA2* may also increase the risk of breast and ovarian cancer.

**Figure 6.**
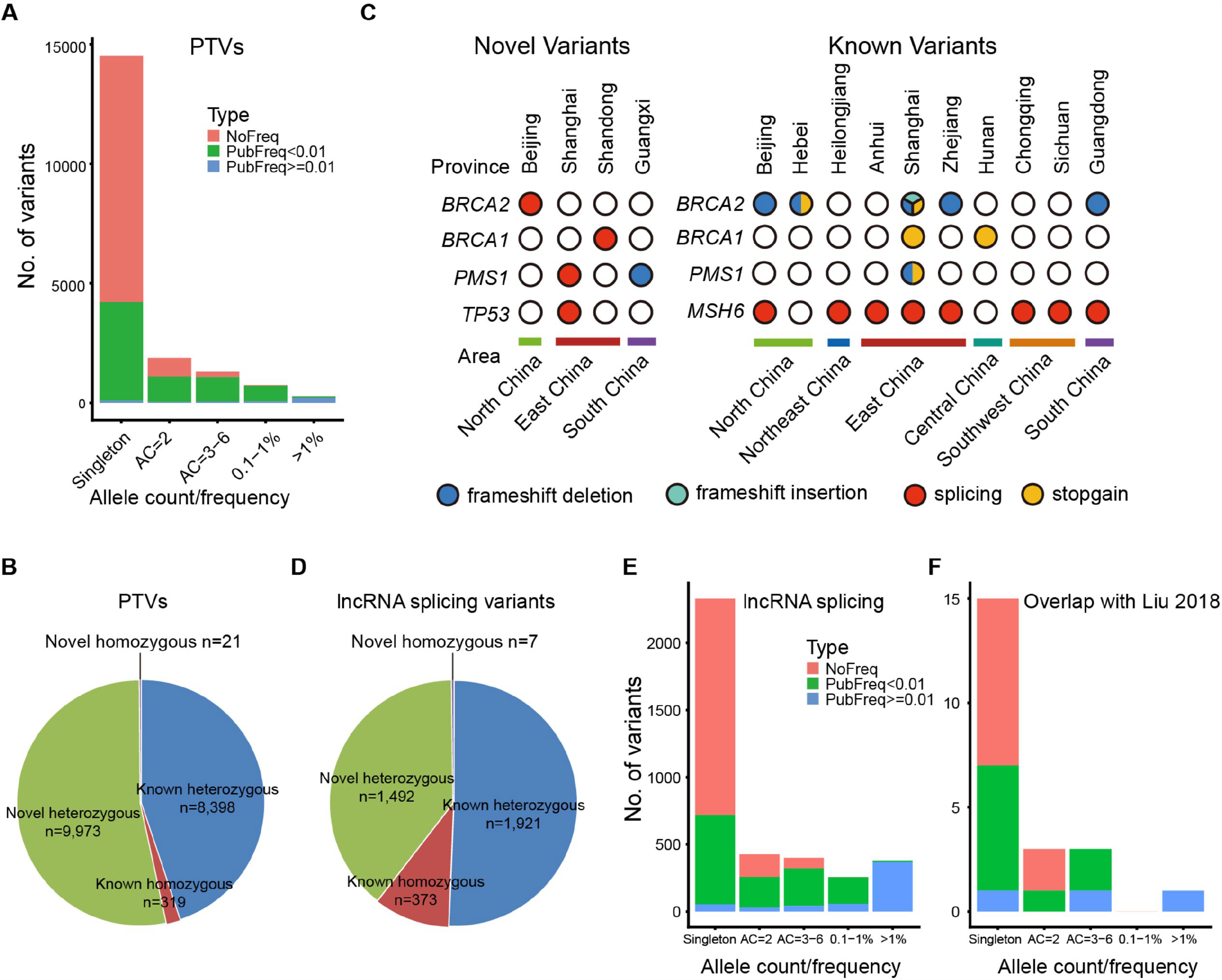
Predicted loss-of-function variants in NyuWa dataset. (A) Allele count and frequency distribution for protein truncating variants (PTVs). (B) Number of protein truncating variants (PTVs) grouped by novel, known, heterozygous and homozygous. (C) Known and novel PTVs in selected cancer associated genes identified in NyuWa dataset. (D) Number of lncRNA splicing variants grouped by novel, known, heterozygous and homozygous. (E) Allele count and frequency distribution for lncRNA splicing variants. (F) Allele count and frequency distribution for lncRNA splicing variants in 230 lncRNA genes reported to be essential for cell growth.

Since lncRNAs do not contain consensus CDS regions, for possible lncRNA loss-of-function variants, splicing variants become the most important class. Splicing variants may cause intron retention or exon skipping, and greatly change the lncRNA sequence and structure (Ulitsky et al. 2011). 230 lncRNA genes were reported to affect cell growth after CRISPR editing at lncRNA splicing sites (Liu et al. 2018b), suggesting the importance of lncRNA splicing variants for lncRNA functions. A total of 3,793 splicing variants in 3,544 lncRNA genes were found in NyuWa dataset (Figure 6D), including 1,454 splicing variants in 1,287 Ensembl lncRNA genes and another 2,339 splicing variants in 2,257 NONCODE lncRNA genes (Supplementary information, Figure S10F and S10G). Each sample had a median of 61 homozygous and 91 heterozygous lncRNA splicing variants (Supplementary information, Figure S10H). Among 230 lncRNA genes reported to be essential for cell growth (Liu et al. 2018b), we found 22 splicing variants in 20 lncRNA genes. The proportion of AF > 0.1% lncRNA splicing variants were relatively smaller in the 20 essential lncRNA genes compared with all lncRNA splicing variants (Figure 6E and 6F), suggesting that splicing variants can really affect functions of these lncRNAs. In general, the loss-of-function variants for both protein-coding and non-coding genes identified in NyuWa dataset may be associated with disease causing or trait tendency, which will provide novel insights into disease and genetic studies.

## Discussion

Chinese population accounts for about 20% of the global human population, with 56 ethnic groups and great diversity of disease types. Constructing a comprehensive genome resource platform of Chinese population empowers medical genetics discoveries in the world’s largest population, and also contributes to the diversity of worldwide human genetic resource. Here we presented the NyuWa resource of large cohort deep WGS data for Chinese population, and constructed a companion database to comprehensively catalogue the variants. The 25.0M novel variants identified in NyuWa resource will greatly benefit studies of human diseases, especially for Chinese people. Although ChinaMAP has also published a resource of Chinese population, the variant data files were not available for downloading. By comparing of manually selected variants, we estimated ∼18M variants would remain novel after subtracting variants in ChinaMAP.

Another important contribution of this work is that the NyuWa resource filled the blank of WGS based haplotype reference panel for Chinese population. Previously, the most commonly used imputation panels were constructed by the 1KGP3 (Auton et al. 2015) and HRC (McCarthy et al. 2016). The recently released TOPMed reference panel included the largest number of haplotypes (Taliun et al. 2019) so far. However, imputation performance of these panels for Chinese and East Asian populations are limited, as East Asian samples are underrepresented. In addition, large number of genome variants are population- or sample-specific, especially for rare variants, imputation of which can be challenging (Carmi et al. 2014). Our NyuWa reference panel contains 19.3M variants (approximately MAF > 0.1%) with 3.25M specific variants not included in other panels, which contains a large proportion of low frequency alleles. The imputation performance of NyuWa outperformed that of 1KGP3, HRC and TOPMed for Chinese population (Figure 3A; Supplementary information, Figure S3A-B). Furthermore, the combined reference panel of NyuWa and 1KGP3 outperformed 1KGP3, HRC and TOPMed for nearly all the Asians (Figure 3A; Supplementary information, Figure S3). Compared to GAsP, a newly public Asian reference panel (Wall et al. 2019), NyuWa also has advantage in Chinese populations including Han, She, Tujia, Miaozu, Yizu, Tu and Naxi, and possesses larger number of high quality imputed variants (Rsq > 0.8) across all MAF bins. However, due to the lack of samples from certain Chinese minority populations, the performance of NyuWa reference panel can still be improved by including more minority samples in the future.

We also found that the genetic differences between north and south Chinese are consistent with differences between two major ADMIXTURE components, suggesting that the north-south differences mainly result from partial population mix in recent history. In the ADMIXTURE results, the difference was mainly the proportion of north Han like component (ancestry 1, red) and south Dai or Vietnamese like component (ancestry 3, blue) (Figure 4A and 4B). The north samples have very large proportion of component 1 and small component 3, while component 3 reaches to about a half in the south samples. This population structure implies a partial mix of two ancestral components of north and south, which is also consistent with the history of China. The earliest center of Chinese civilization located in the central to north of China, ranging from Henan to Shaanxi. Starting from the Eastern Zhou Dynasty, the Chinese territory expanded greatly, especially to the south. Then the foundation of unitary multiethnic country beginning at Qin and Han Dynasty facilitated the mix of early Chinese population with south ancestral populations. The mix has still not achieved equilibrium up to now. Despite the lack of native place identities for many samples in NyuWa resource, we could still detect a clear difference between north and south Chinese samples, indicating that the hospitals collecting these samples were good approximations.

An ideal reference panel for a population needs to cover all major ADMIXTURE components in the population. Each major component is required to have sufficient and balanced sample size to cover most haplotypes in the component. As described above, both north and south Han Chinese consist of the same two major components, though the proportions of these components are different. So the same reference panel that cover these major components can be used to impute both north and south Han populations. Imputation tests using north or south subset panels confirmed the speculation. These results are based on the current sample size. In future when sample size is large enough, which panel works better still depends on the specific situation.

The current knowledge and guidelines about medical genomics are mainly from Eurocentric genetic and genomic resources, and may miss information about non-European ancestry. Our study provides a large and high-quality WGS resource for Chinese populations, which will help to examine the effect of known genetic variants on disease susceptibility and drug responses, and benefit clinical investigations in the future. The identification of loss-of-function variants for both protein-coding and lncRNA genes in this study expands the catalogue of loss-of-function variants in nature. When combined with phenotype information, this resource will provide important biological insights into gene functions.

## Methods

### DNA extraction and library preparation

Genomic DNA was extracted and sequenced by WuXi Apptec Co., Ltd. according to the standard protocols of Illumina on HiSeq X10 platform or NovaSeq 6000. The sequencing reads were paired-end 150 nt and the target depth is 30X. Sequencing quality was checked with FastQC v0.11.3 (http://www.bioinformatics.babraham.ac.uk/projects/fastqc). Adaptor sequences and low quality bases were removed with Trimmomatic v0.36 (Bolger et al. 2014).

### Variant calling

The variant calling pipeline followed GATK (Ryan Poplin 2017) Best Practices Workflows Germline short variant discovery (SNPs + Indels) joint genotyping cohort mode. In brief, the raw sequencing reads were mapped to human reference genome assembly 38 with BWA-MEM v0.7.15 (Li and Durbin 2010). Picard (http://broadinstitute.github.io/picard/) was used to sort bam and mark duplicates. Mapping quality was check by qualimap v2.1.2 (Okonechnikov et al. 2016). Indels were realigned and bases were recalibrated with GATK v3.7. Variants were called for each sample using GATK HaplotypeCaller in ‘GVCF’ mode. GATK GenotypeGVCFs was then used to identify variants for all samples in the cohort. Then GATK VQSR was applied for SNPs and indels with truth sensitivity filter levels 99.7 and 99.0, respectively. Variants were then annotated with annovar v2018-04-16 (Wang et al. 2010).

### Sample and site filtering

Duplicate sequencing data for same persons were removed. verifyBamID2 (Zhang et al. 2020) version 1.0.6 was used to check the contamination. Samples with contamination level alpha >= 0.05 were removed. Sex of each sample was inferred by two ways. Based on whole genome and chromosome coverage results reported by qualimap, the coverage of X and Y chromosomes were divided by the whole genome coverage. The relative coverage of (X, Y) of male is expected to be (0,5, 0.5), and that of female is expected to be (1, 0). The ploidy of non-PAR region of X chromosome were estimated by BCFtools v1.5 (https://samtools.github.io/bcftools/bcftools.html) guess-ploidy module. Males are haploid while females are diploid.

To filter low quality sites, variants with VQSR not passed were removed. Additional filters were applied to further exclude low quality variants. Sites with genotype quality (GQ) < 10 in > 50% samples were removed. For Y chromosome, sites were removed if GQ < 10 in > 50% male samples, or GQ >= 10 in > 10% female samples. Sites with no ALT allele in GQ >= 10 samples were also removed. Variants were further filtered with a Hardy-Weinberg Equilibrium (HWE) p value < 10^−6^ in the direction of excessive heterozygosity or ExcessHet > 54.69 in the INFO column calculated by GATK. Multi-allele sites were split using BCFtools norm module.

Some analyses required removal of close relatives. 3^rd^ degree or closer relationships were identified with the combination of kinship coefficient (Φ) and probability of zero identity-by-descent (IBD) sharing (π_0_) (Manichaikul et al. 2010) calculated by plink (Chang et al. 2015). The k-degree relationship was defined as 2^-k-1.5^ < Φ < 2^-k-0.5^. For the 1^st^ degree relationships, parent-offspring was defined as π _0_ < 0.1 and full sibling if π _0_ > 0.1. Φ> 2^-1.5^ represents monozygotic twin or sample replicates. Relationships more than 3^rd^ degree were treated as unrelated. To determine the list of excluded close relatives, samples with more relatives were excluded with priority, and a maximum of 2,902 unrelated samples were kept.

### Haplotype phasing

Sequencing reads based haplotype phasing for each sample was carried out with HAPCUT2 (Edge et al. 2017). The local phased sets were then incorporated in population-based phasing of 2,999 samples using SHAPEIT4 (Delaneau et al. 2019) version 4.1.2 with parameter ‘--use-PS 0.0001’. The information from family trios or duos were converted to phasing scaffold data and used by SHAPEIT4 with ‘--scaffold’ option. Sites with missing call rate greater than 10% were removed. Sites with minor allele count < 2 (MAC2) were also removed. There were no samples with missing call rate greater than 10%. No additional reference panel was used. Only chromosome 1-22 and X were phased and each chromosome was phased separately. For X chromosome, the pseudo-autosomal regions (PARs) and non-PAR were divided and phased separately. For samples with haploid X chromosome in non-PAR regions (male), the heterozygous genotypes were converted to missing before phasing using SHAPEIT4.

### Reference panel

The 2,902 independent samples were extracted from phased data above. Sites with minor allele count < 5 (MAC5) in the independent sample set were also removed. The final list included 2,902 samples and 19,256,267 variants. Phased genotypes were then converted to m3vcf format as imputation reference file using Minimac3 (Das et al. 2016) v2.0.1. The hg38 version of 1KGP3 reference panel was generated similarly with MAC5 sites.

To further improve imputation performance, a combined panel of NyuWa with 1KGP3 panel was generated using the reciprocal imputation strategy (Huang et al. 2015). The missed variants in each panel were imputed with the other with Minimac4 (Das et al. 2016), and the results were combined to form a new panel with all samples and union of variants in NyuWa and 1KGP3 panel. The combined panel had 40,196,029 variants in total.

### Imputation performance

The chromosome 2 of HGDP data was used to test imputation performance for NyuWa, 1KGP3, GAsP, HRC.r1.1, TOPMed and NyuWa+1KGP3 reference panels. Bi-allele SNPs that exist in all panels were selected. Then every 1 out of 10 of the selected SNPs were masked to evaluate the imputation accuracy. Phasing and imputation of GAsP HRC.r1.1 and TOPMed panel were run on respective web servers. Phasing and imputation of NyuWa, 1KGP3 and NyuWa+1KGP3 panels were run locally with Eagle2 (Loh et al. 2016) and Minimac4, respectively. Imputation error rate was computed for each population as the genotype discordance rate of the masked SNPs. In addition, for Chinese and Han Chinese samples in HGDP dataset, we compared the Rsq statistic for total imputed variants in different MAF bins (MAF: <=0.01, 0.01-0.05, 0.05-0.2, and 0.2-0.5).

The imputation error rates of reference panels constructed with sample subsets of NyuWa reference panel were evaluated the same way as NyuWa panel. The 1KGP3 CHS and CHB test samples were already phased, and every 1 out of 10 of the selected SNPs were masked to evaluate the imputation error rates. The samples in the North or South specific panels were divided based on ranks of sample positions on PC1 from PCA or geographical demarcation of Qinling Mountains-Huaihe River (Supplementary information, Table S1).

### Population structure analysis

NyuWa 2,902 independent samples and 1KGP3 data were merged by extracting overlapped bi-allelic autosomal SNPs. SNPs with missing rate of more than 10% or MAF less than 0.05 were excluded. Linkage equilibrium (LD) was removed by thinning the SNPs to no closer than 2kb using plink. Furthermore, 27 known long-range LD regions were removed according to previous studies (Price et al. 2008; Tang et al. 2008; Wu et al. 2019). The resulted dataset included 901,455 SNPs. The merged data were then used in principal component analysis (PCA) and ADMIXTURE by extracting samples of interest in each analysis. PCA was carried out using plink. ADMIXTURE were carried out using ADMIXTURE Version 1.3.0 (Alexander et al. 2009). For each K, the analysis was repeated 4 to 8 times with different seeds, and the one with the highest value of likelihood was chosen. For ADMIXTURE result display when K > 2, dimensions were reduced to 1-dimension by tSNE and samples were ordered by tSNE values.

### F_st_ between south and north of China

SNP-level fixation index (F_st_) between north and south of China was calculated using the Weir and Cockerham’s estimator (Weir and Cockerham 1984) integrated in VCFtools (Danecek et al. 2011). North and south of China were divided according to the classic demarcation of Qinling Mountains-Huaihe River (Supplementary information, Table S1). Henan, Jiangsu, Anhui were excluded because the Huaihe River flows through these provinces. Shanghai was also excluded for the possibility that there may be too many individuals from other provinces.

### Introgression of Denisovan and Neanderthal ancestry

Estimation of Denisovan and Neanderthal ancestry followed methods in GAsP (Wall et al. 2019). In brief, Neanderthal and Denisovan genomes were downloaded from http://cdna.eva.mpg.de/neandertal/altai/AltaiNeandertal/VCF/ and http://cdna.eva.mpg.de/neandertal/altai/Denisovan/. Human ancestral sequences were downloaded from ftp://ftp.ensembl.org/pub/release-99/fasta/ancestral_alleles/. Potential Neanderthal/Denisovan SNPs were filtered by the following criteria. 1. The REF allele matched the ancestral allele; 2. Neanderthal/Denisovan genotype was homozygous ALT allele; 3. Denisovan/Neanderthal genotype was homozygous REF allele; 4. ALT allele was not found in YRI, GWD, MSL or ESN samples in 1KGP3. Then, for each NyuWa sample, the number of Neanderthal/Denisovan SNP alleles were counted. To correct background, linear models were fit for both Neanderthal and Denisovan SNPs based on allele counts and ancestry percentage in GAsP results. Supposing SNPs called in NyuWa and GAsP were independent for Neanderthal/Denisovan SNPs, allele counts were scaled to make the median of NyuWa samples equal to the average of GAsP HAN samples. The ancestry proportion for each sample was then determined by the linear model using scaled allele count.

### Y chromosome analysis

Genotypes of male chrY SNPs in NyuWa dataset were lift over to hg19 using CrossMap (Zhao et al. 2014). Y chromosomal haplogroups were inferred using yHaplo (https://github.com/23andMe/yhaplo) (Poznik 2016). Besides, file of primary tree structure (y.tree.primary.2016.01.04.nwk), file of preferred SNP names (preferred.snpNames.txt) and file of phylogenetically informative SNPs (isogg.2016.01.04.txt) were used.

MEGA X (Kumar et al. 2018) were used to construct a phylogenetic tree based on neighbor joining (NJ) method with 50 bootstrap. FigTree v1.4.4 (https://github.com/rambaut/figtree/releases) was used to colour the tree and label main branches manually.

### Protein-truncating variants (PTVs) and lncRNA loss-of-function splicing variants

PTV analysis followed methods in GAsP (Wall et al. 2019). In brief, stop gain, frameshift and splicing sites were selected according to ensGene annotation by annovar (Wang et al. 2010). Splicing variants are variants within 2-bp away from an exon/intron boundary that disrupt the GT-AG boundary pattern. Then multiple filters were applied. Variants out of Genome In A Bottle (GIAB) high confidence regions were excluded. Stop gain or frameshift variants in the last exon or the last 50 nt in the second last exon were excluded. Variants in exons with non-classic splice sites were also removed. Splicing variants that locate in introns length < 15 nt or UTRs were excluded. Stop gain and splicing variants with phyloP100way vertebrate rankscore < 0.01 were excluded. Additional filters were applied to filter high quality PTVs. Only variants with GQ >= 20, DP > 7 and ALT DP > DPΦ0.2 were kept. Only variants affecting transcripts that within top 50% of gene expression in GTEx (Ardlie et al. 2015) were kept. A total of 9,526 PTVs in 4666 genes were obtained.

Loss-of-function variants were also predicted using LOFTEE v 1.0.3 (https://github.com/konradjk/loftee). A total of 16,910 High confidence loss-of-function variants in canonical transcripts were identified. These variants covered most (7,725) of previously identified PTVs. The results were then combined to get the union set of PTVs.

For lncRNA splicing variants, Ensembl annotation was used first. Splicing variants were filtered similar to PTVs except that the phyloP100way conservation filter was not applied. The remaining splicing variants in NONCODE annotation were also filtered similarly, with GTEx expression replaced with expression data downloaded from NONCODE database.

## Data access

The datasets generated and/or analysed during the current study are available at http://bigdata.ibp.ac.cn/NyuWa/.

## Acknowledgments

We thank Weiwei Zhai for thoughtful discussions and valuable comments on the population structure analysis.

## Author Contributions

T.X. and S.H. conceptualized and supervised the project. P.Z., H.L., Y.L., J.W., Y.N., Q.K., Y.S. and H.Z. conducted analysis. Y.W. and T.X. contributed to sample collection and data generation. P.Z., H.L., Y.Z., Q.K. and T.S. made the web server and database. P.Z., H.L., Y.N., and S.H. drafted the manuscript, and all the primary authors reviewed, edited, and approved manuscript.

