## Supplementary Figures and Tables for "NyuWa Genome Resource: Deep Whole Genome Sequencing Based Chinese Population Variation Profile and Reference Panel"

Table S1. Summary of samples in different provinces and regions.

| Region | Province | No. of samples | North/South for Fst |
|---|---|---|---|
| Northeast China | Heilongjiang | 39 | North |
| | Jilin | 8 | North |
| | Liaoning | 2 | North |
| North China | Beijing | 305 | North |
| | Inner Mongolia | 6 | North |
| | Tianjing | 10 | North |
| | Hebei | 42 | North |
| | Shanxi | - | - |
| Northwest China | Shaanxi | 31 | North |
| | Xinjiang | 1 | North |
| | Gansu | - | - |
| | Ningxia | - | - |
| | Qinghai | - | - |
| East China | Shanghai | 1822 | - |
| | Shandong | 15 | North |
| | Jiangsu | 17 | - |
| | Anhui | 89 | - |
| | Zhejiang | 124 | South |
| | Jiangxi | 9 | South |
| | Fujian | 1 | South |
| | Taiwan | - | - |
| Central China | Henan | 2 | - |
| | Hubei | 41 | South |
| | Hunan | 23 | South |
| South China | Guangdong | 329 | South |
| | Guangxi | 15 | South |
| | Hainan | - | - |
| | HongKong | - | - |
| | Macao | - | - |
| Southwest China | Sichuan | 45 | South |
| | Chongqing | 23 | South |
| | Guizhou | - | - |
| | Yunnan | - | - |
| | Tibet | - | - |

Table S2. Number of variants detected in different chromosomes

| type | SNP | | | insertion | | | deletion | | |
|---|---|---|---|---|---|---|---|---|---|
| **CHR** | **novel** | **total** | **novel pecentage** | **novel** | **total** | **novel pecentage** | **novel** | **total** | **novel pecentage** |
| **chr1** | 1,786,646 | 5,457,328 | 32.74% | 46,960 | 260,688 | 18.01% | 102,619 | 371,920 | 27.59% |
| **chr2** | 1,982,921 | 5,979,062 | 33.16% | 51,288 | 274,940 | 18.65% | 113,605 | 405,812 | 27.99% |
| **chr3** | 1,654,532 | 4,947,700 | 33.44% | 41,790 | 230,006 | 18.17% | 94,241 | 333,998 | 28.22% |
| **chr4** | 1,574,474 | 4,755,496 | 33.11% | 39,743 | 224,053 | 17.74% | 91,298 | 329,067 | 27.74% |
| **chr5** | 1,479,739 | 4,445,567 | 33.29% | 37,422 | 203,466 | 18.39% | 84,673 | 299,956 | 28.23% |
| **chr6** | 1,370,917 | 4,258,187 | 32.19% | 36,150 | 206,514 | 17.50% | 80,791 | 298,726 | 27.05% |
| **chr7** | 1,284,098 | 3,976,486 | 32.29% | 33,476 | 187,464 | 17.86% | 74,103 | 273,212 | 27.12% |
| **chr8** | 1,265,796 | 3,854,220 | 32.84% | 30,766 | 166,648 | 18.46% | 68,471 | 243,252 | 28.15% |
| **chr9** | 973,092 | 3,019,272 | 32.23% | 24,915 | 135,254 | 18.42% | 53,446 | 192,997 | 27.69% |
| **chr10** | 1,076,252 | 3,374,044 | 31.90% | 28,227 | 157,610 | 17.91% | 61,315 | 225,894 | 27.14% |
| **chr11** | 1,101,638 | 3,381,635 | 32.58% | 27,955 | 151,727 | 18.42% | 62,664 | 221,056 | 28.35% |
| **chr12** | 1,058,196 | 3,243,039 | 32.63% | 28,543 | 160,813 | 17.75% | 62,658 | 230,478 | 27.19% |
| **chr13** | 797,179 | 2,415,877 | 33.00% | 20,984 | 117,314 | 17.89% | 47,412 | 172,294 | 27.52% |
| **chr14** | 719,140 | 2,212,627 | 32.50% | 19,167 | 105,904 | 18.10% | 41,867 | 152,770 | 27.41% |
| **chr15** | 650,804 | 2,028,103 | 32.09% | 17,961 | 96,139 | 18.68% | 37,489 | 135,494 | 27.67% |
| **chr16** | 716,067 | 2,310,781 | 30.99% | 17,531 | 97,639 | 17.95% | 36,455 | 138,134 | 26.39% |
| **chr17** | 614,070 | 1,983,616 | 30.96% | 18,015 | 101,398 | 17.77% | 36,111 | 139,765 | 25.84% |
| **chr18** | 618,010 | 1,897,311 | 32.57% | 16,233 | 88,991 | 18.24% | 36,052 | 130,533 | 27.62% |
| **chr19** | 453,971 | 1,572,671 | 28.87% | 13,237 | 83,833 | 15.79% | 26,381 | 111,815 | 23.59% |
| **chr20** | 504,689 | 1,584,965 | 31.84% | 13,488 | 72,961 | 18.49% | 28,292 | 104,304 | 27.12% |
| **chr21** | 277,236 | 880,444 | 31.49% | 7,480 | 43,557 | 17.17% | 16,527 | 63,288 | 26.11% |
| **chr22** | 276,589 | 946,066 | 29.24% | 8,136 | 45,235 | 17.99% | 15,929 | 63,312 | 25.16% |
| **chrX** | 876,878 | 2,525,483 | 34.72% | 14,890 | 140,858 | 10.57% | 31,765 | 171,538 | 18.52% |
| **chrY** | 2,444 | 7,981 | 30.62% | 223 | 2,953 | 7.55% | 524 | 2,810 | 18.65% |
| **ALL** | 23,115,378 | 71,057,961 | 32.53% | 594,580 | 3,355,965 | 17.72% | 1,304,688 | 4,812,425 | 27.11% |

Table S3. Allele frequency and genomic features of the SNVs and indels annotated by RefSeq

| type | AF | exonic | splicing | intronic | UTR5 | UTR3 | ncRNA_exonic | ncRNA_splicing | ncRNA_intronic | upstream | downstream | intergenic | TOTAL |
|---|---|---|---|---|---|---|---|---|---|---|---|---|---|
| SNP | af<=0.001 | 701,486 | 6,213 | 21,226,561 | 141,082 | 643,596 | 222,761 | 1,263 | 3,428,531 | 389,235 | 376,729 | 28,727,452 | 55,864,909 |
| | 0.001-0.01 | 71,583 | 380 | 2,571,965 | 15,481 | 74,885 | 28,029 | 161 | 423,561 | 46,865 | 47,229 | 3,610,650 | 6,890,789 |
| | 0.01-0.05 | 16,981 | 90 | 758,557 | 4,167 | 21,144 | 8,240 | 64 | 129,721 | 13,834 | 14,185 | 1,114,504 | 2,081,487 |
| | 0.05-0.5 | 25,588 | 101 | 1,480,399 | 6,886 | 37,056 | 16,001 | 95 | 261,681 | 26,157 | 26,761 | 2,265,539 | 4,146,264 |
| | 0.5-1 | 12,097 | 59 | 724,412 | 3,433 | 18,497 | 7,863 | 43 | 131,611 | 13,199 | 13,498 | 1,149,800 | 2,074,512 |
| | ALL | 827,735 | 6,843 | 26,761,894 | 171,049 | 795,178 | 282,894 | 1,626 | 4,375,105 | 489,290 | 478,402 | 36,867,945 | 71,057,961 |
| Indel | af<=0.001 | 24,695 | 1,548 | 1,936,879 | 11,964 | 77,258 | 17,040 | 111 | 310,919 | 37,574 | 40,149 | 2,449,642 | 4,907,779 |
| | 0.001-0.01 | 2,757 | 105 | 599,015 | 2,871 | 18,897 | 3,751 | 22 | 95,227 | 11,263 | 11,609 | 778,182 | 1,523,699 |
| | 0.01-0.05 | 615 | 34 | 269,718 | 898 | 7,511 | 1,408 | 13 | 41,628 | 5,026 | 5,290 | 350,977 | 683,118 |
| | 0.05-0.5 | 640 | 48 | 289,888 | 979 | 8,184 | 1,885 | 13 | 50,022 | 5,774 | 5,984 | 422,034 | 785,451 |
| | 0.5-1 | 216 | 24 | 98,282 | 453 | 3,050 | 756 | 3 | 16,937 | 1,903 | 2,034 | 144,685 | 268,343 |
| | ALL | 28,923 | 1,759 | 3,193,782 | 17,165 | 114,900 | 24,840 | 162 | 514,733 | 61,540 | 65,066 | 4,145,520 | 8,168,390 |

Table S4. Allele frequency and genomic features of the SNPs and indels annotated by NONCODE V5

| type | AF | ncRNA_exonic | ncRNA_splicing | ncRNA_intronic | upstream | downstream | intergenic | TOTAL |
|---|---|---|---|---|---|---|---|---|
| **SNP** | **af<=0.001** | 3,413,380 | 10,982 | 16,330,425 | 1,218,737 | 1,082,596 | 33,808,789 | 55,864,909 |
| | **0.001-0.01** | 417,088 | 1,302 | 2,006,382 | 150,814 | 135,501 | 4,179,702 | 6,890,789 |
| | **0.01-0.05** | 124,448 | 426 | 611,884 | 45,297 | 41,195 | 1,258,237 | 2,081,487 |
| | **0.05-0.5** | 246,502 | 896 | 1,237,712 | 88,814 | 79,987 | 2,492,353 | 4,146,264 |
| | **0.5-1** | 122,739 | 451 | 626,395 | 44,394 | 39,821 | 1,240,712 | 2,074,512 |
| | **ALL** | 4,324,157 | 14,057 | 20,812,798 | 1,548,056 | 1,379,100 | 42,979,793 | 71,057,961 |
| **Indel** | **af<=0.001** | 290,471 | 1,178 | 1,466,459 | 113,366 | 102,100 | 2,934,205 | 4,907,779 |
| | **0.001-0.01** | 78,130 | 339 | 451,918 | 36,285 | 33,670 | 923,357 | 1,523,699 |
| | **0.01-0.05** | 33,169 | 150 | 200,843 | 16,317 | 15,344 | 417,295 | 683,118 |
| | **0.05-0.5** | 39,981 | 187 | 236,529 | 18,284 | 17,072 | 473,398 | 785,451 |
| | **0.5-1** | 14,019 | 84 | 81,749 | 6,401 | 5,758 | 160,332 | 268,343 |
| | **ALL** | 455,770 | 1,938 | 2,437,498 | 190,653 | 173,944 | 4,908,587 | 8,168,390 |

Table S5. Statistics of SNPs and indels in coding regions

| type | AF | frameshift deletion | frameshift insertion | nonframeshift deletion | nonframeshift insertion | nonsynonymous SNV | synonymous SNV | stopgain | stoploss | unknown | TOTAL |
|---|---|---|---|---|---|---|---|---|---|---|---|
| **SNV** | **af<=0.001** | NA | NA | NA | NA | 433,851 | 254,132 | 10,431 | 437 | 2,856 | 701,707 |
| | **0.001-0.01** | NA | NA | NA | NA | 40,515 | 30,025 | 640 | 36 | 392 | 71,608 |
| | **0.01-0.05** | NA | NA | NA | NA | 8,919 | 7,784 | 104 | 9 | 172 | 16,988 |
| | **0.05-0.5** | NA | NA | NA | NA | 12,106 | 13,135 | 119 | 12 | 226 | 25,598 |
| | **0.5-1** | NA | NA | NA | NA | 5,575 | 6,374 | 29 | 6 | 122 | 12,106 |
| | **ALL** | NA | NA | NA | NA | 500,966 | 311,450 | 11,323 | 500 | 3,768 | 828,007 |
| | | | | | | | | | | | 0 |
| **Indel** | **af<=0.001** | 9,929 | 3,946 | 6,973 | 3,080 | NA | NA | 550 | 95 | 148 | 24,721 |
| | **0.001-0.01** | 669 | 345 | 1,116 | 559 | NA | NA | 46 | 11 | 14 | 2,760 |
| | **0.01-0.05** | 126 | 80 | 220 | 165 | NA | NA | 9 | 5 | 10 | 615 |
| | **0.05-0.5** | 115 | 78 | 236 | 195 | NA | NA | 7 | 2 | 9 | 642 |
| | **0.5-1** | 37 | 22 | 81 | 74 | NA | NA | 2 | 0 | 6 | 222 |
| | **ALL** | 10,876 | 4,471 | 8,626 | 4,073 | NA | NA | 614 | 113 | 187 | 28,960 |

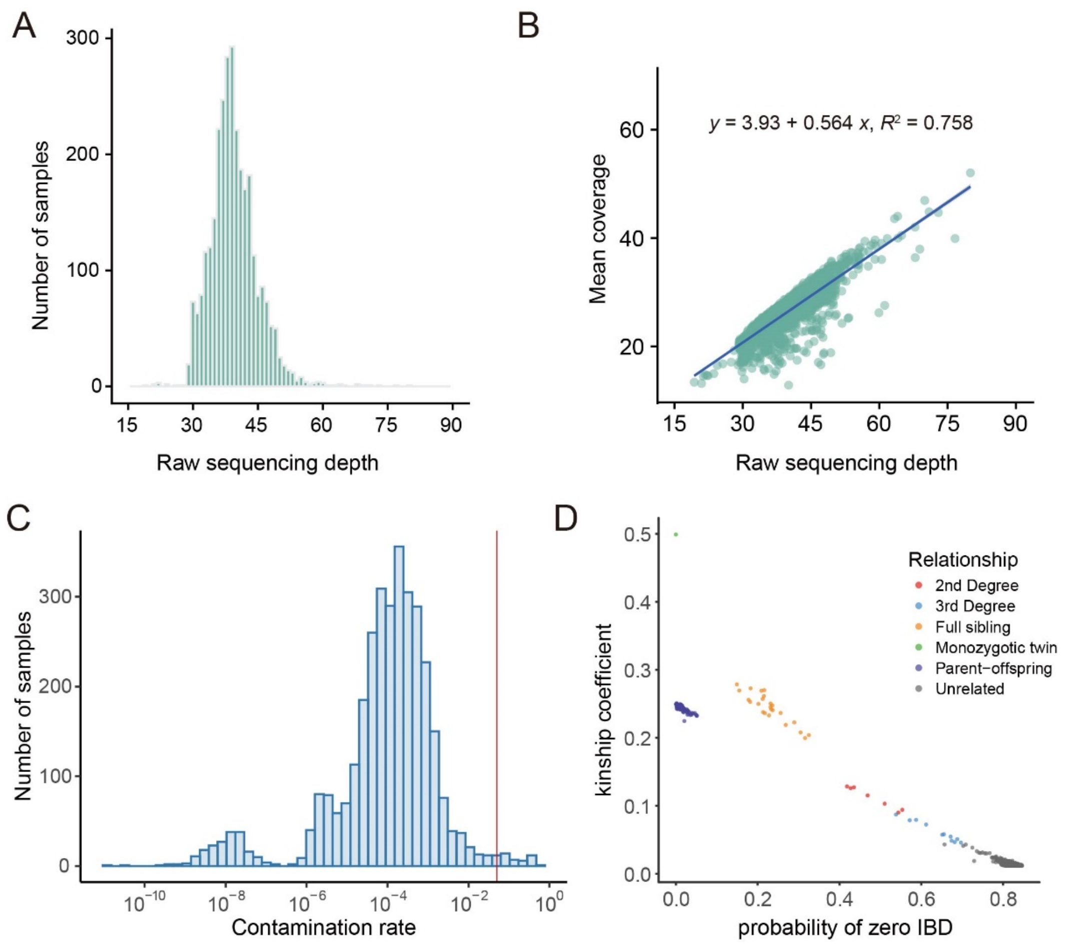


Figure S1. Quality and sample overview.

(A) Distribution of raw sequencing depth in NyuWa resource.

(B) The relationship of WGS raw sequencing depth and mean genomic coverage after removal of duplicates.

(C) Distribution of contamination rates estimated by verifyBamID. A cutoff of 0.05 was used to filter contaminated samples.

(D) Inferred genetic relationship based on kinship coefficient and probability of zero IBD for pairs of individuals.

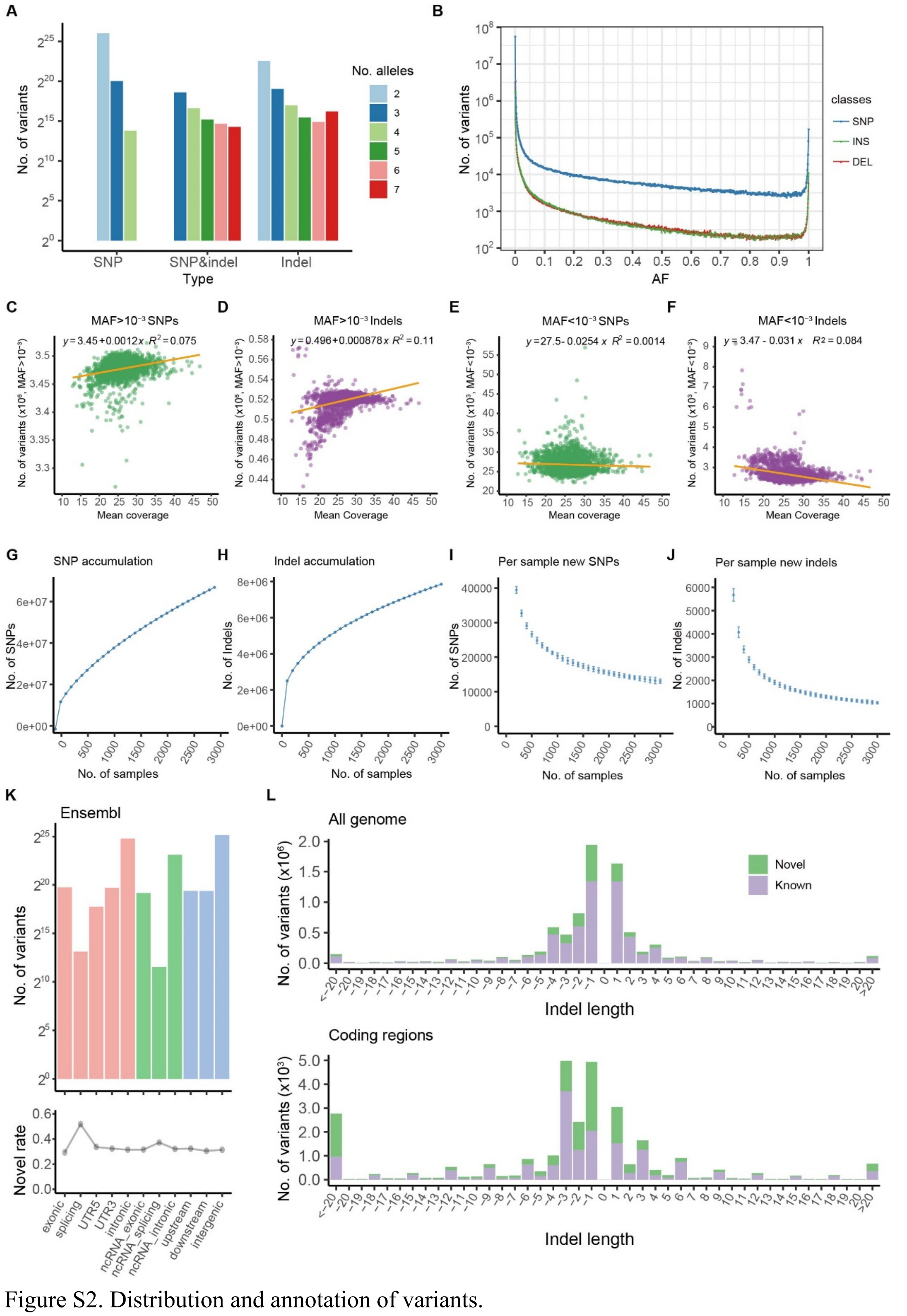

Figure S2. Distribution and annotation of variants.

(A) Distribution of sites with different number of multi-alleles in NyuWa dataset.

(B) Number of variants in different allele frequencies. Each point represents variants in a bin of 0.1% AF.

(C-F) Relationship of genomic coverage (sequencing depth) and number of SNPs (C, E) or indels (D, F) with MAF > $10^{-3}$ (C, D) or MAF < $10^{-3}$ (E, F) detected in each sample.

(G-J) Accumulation of SNPs and indels estimated by randomly downsampling 20 times in NyuWa dataset by intervals of 100 samples. (G) Number of SNPs detected related to sample size. (H) Number of indels detected related to sample size. (I) Number of new SNPs detected by adding one new sample for different sample sizes. Points are shown as mean±SD. (J) Number of new indels detected by adding one new sample for different sample sizes. Points are shown as mean ± SD.

(K) Number of variants (upper) and rate of novel variants (lower) in different Ensembl annotation regions.

(L) Number of variants with different indel lengths in all genomic regions (up) and coding regions (down).

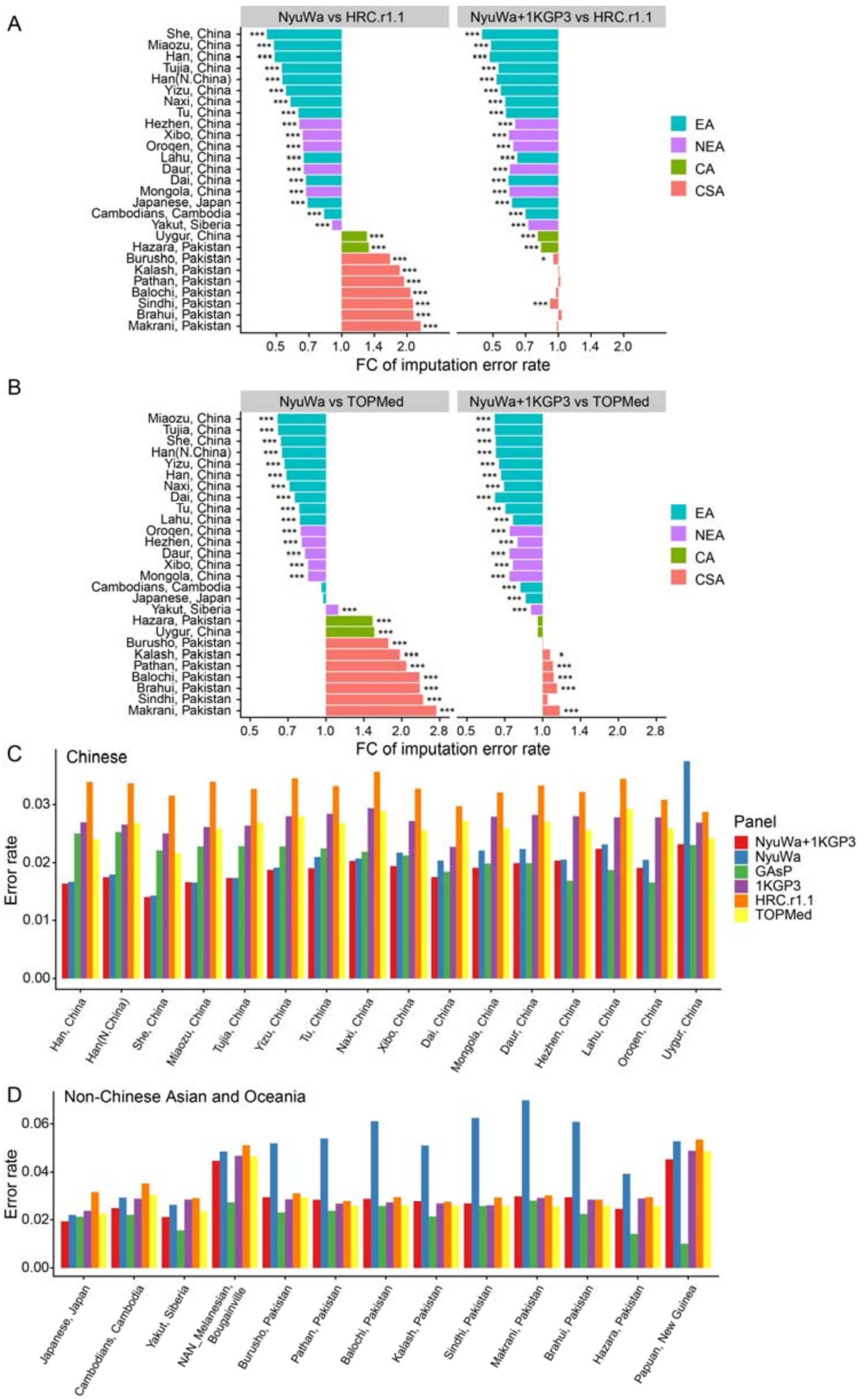


Figure S3. Imputation performance.

(A) Fold change of imputation error rate in different Asia populations in HGDP dataset between HRC.r1.1 panel and NyuWa (left) or NyuWa+1KGP3 (right) panel. EA: East Asian; NEA: North East Asian; CA: Central Asian; CSA: Central South Asian. Significances of error rate differences were calculated by chi-squared test. *: $p < 0.05$; **: $p < 0.01$; ***: $p < 0.001$.

(B) Fold change of imputation error rate in different Asia populations in HGDP dataset between

TOPMed r2 panel and NyuWa (left) or NyuWa+1KGP3 (right) panel. Colors representing regions in (A) and (B) are consistent.

(C) Imputation error rates in different Chinese populations by different reference panels.

(D) Imputation error rates in different non-Chinese Asian and Oceania populations by different reference panels. Colors representing reference panels in (C) and (D) are consistent.

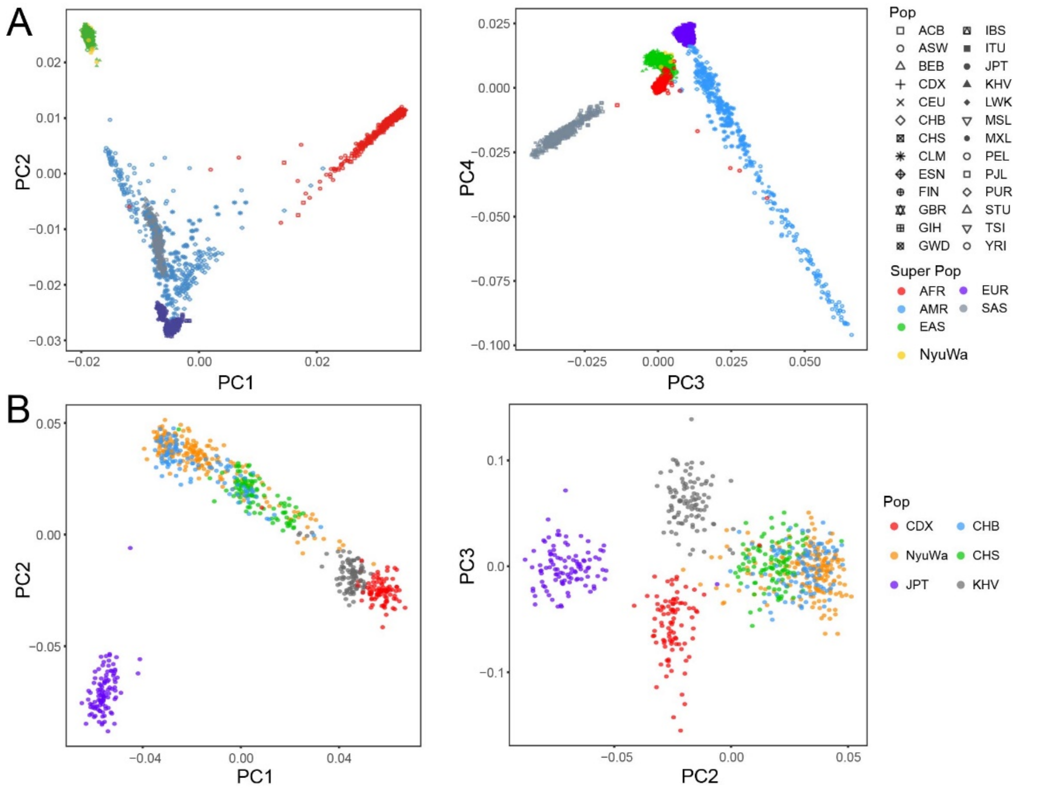


Figure S4. Population structure.

(A) PCA of all 1KGP3 samples with 200 randomly selected NyuWa samples. Abbreviations of populations are from 1KGP3.

(B) PCA of East Asia samples in 1KGP3 with 200 randomly selected NyuWa samples. CHB: Han Chinese in Beijing, China; CHS: Southern Han Chinese; CDX: Chinese Dai in Xishuangbanna, China; JPT: Japanese in Tokyo, Japan; KHV: Kinh in Ho Chi Minh City, Vietnam.

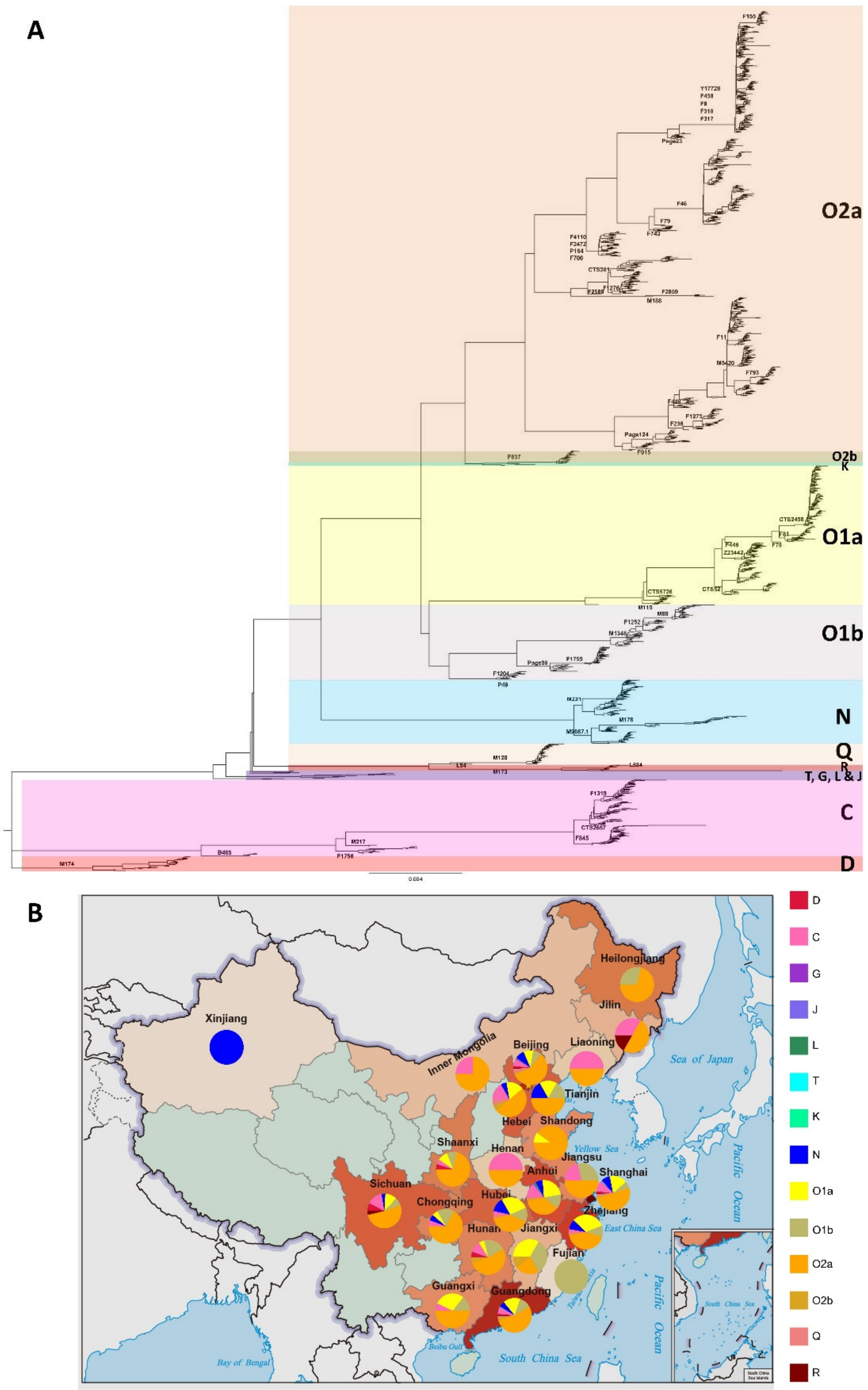


Figure S5. Y haplogroup analysis.

(A) Phylogenetic tree of NyuWa male samples. Y haplogroups are marked with different colors.

(B) Proportions of Y haplogroups in different provinces.

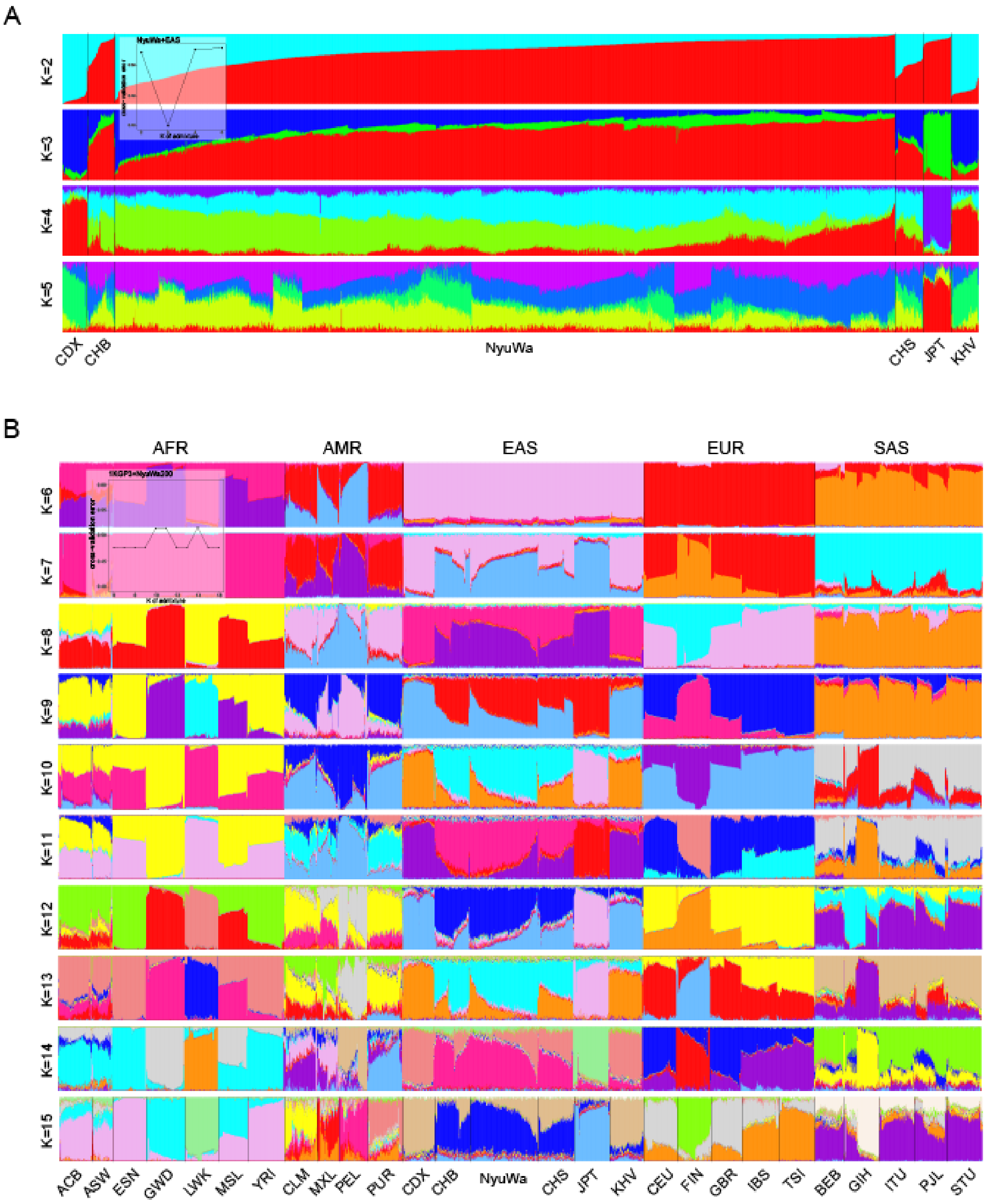


Figure S6. ADMIXTURE analysis of NyuWa and 1KGP3 samples.

(A) ADMIXTURE analysis of all NyuWa samples with 1KGP3 East Asia samples for different Ks. 5-fold cross validation errors were shown. Abbreviations of populations are from 1KGP3.

(B) ADMIXTURE analysis of all 1KGP3 samples and 200 randomly selected NyuWa samples for different Ks. Abbreviations of populations are from 1KGP3.

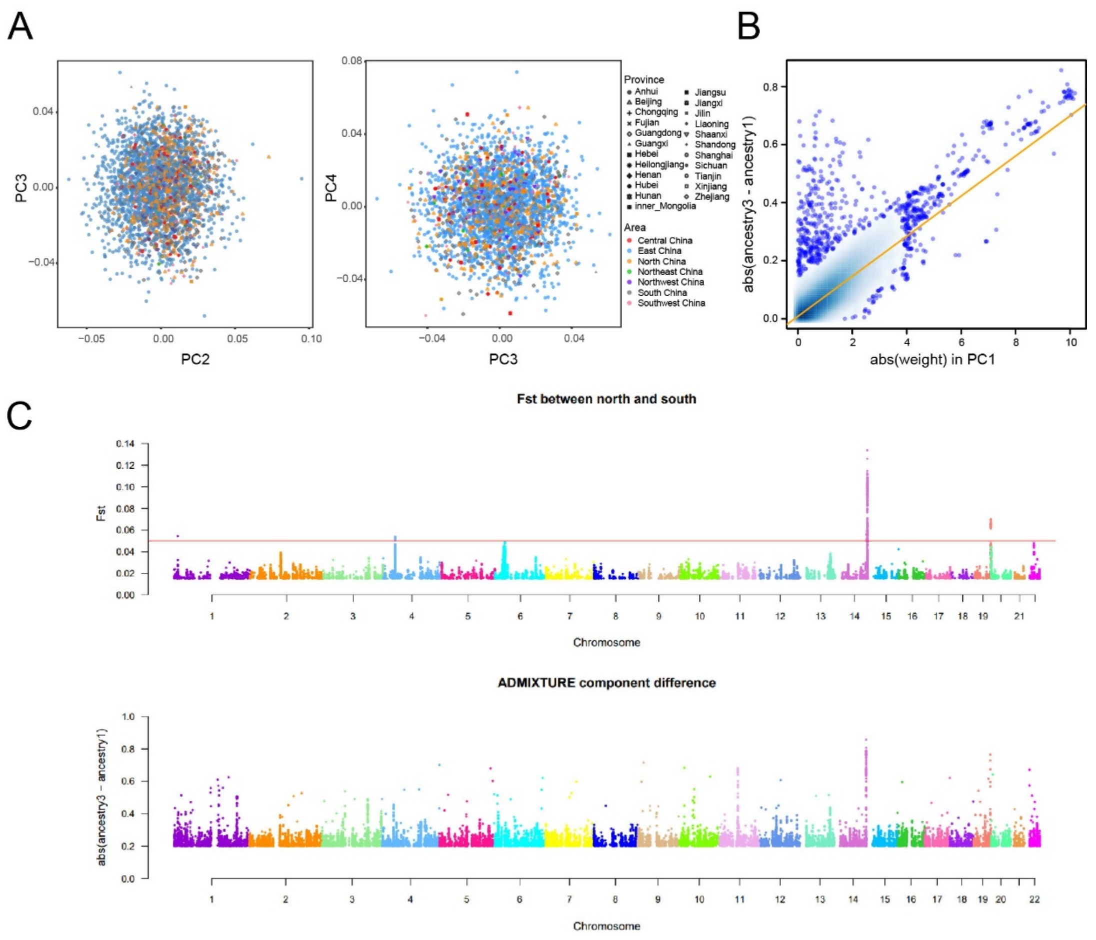


Figure S7. Genetic differences between north and south Han Chinese.

(A) Primary components 2-4 of NyuWa samples. Samples were marked by provinces and areas of China.

(B) Correlation between variant weights in PC1 in Figure 4C (X axis) and differences of allele frequencies in the two major ancestry components in Figure 4A (Y axis). Each point represents a variant locus. The variant AFs in ancestry components were calculated by ADMIXTURE. Variants that contribute greatly to PC1 also show difference of AFs in the two ancestry components.

(C) $F_{st}$ matches differences between ancestry 1 and ancestry 3. SNP level $F_{st}$ scores between north and south samples (upper). North and South of China were divided according to the classic demarcation of Qinling Mountains-Huaihe River (Table S1). Henan, Jiangsu, Anhui were excluded because the Huaihe River flows through these provinces. Shanghai was also excluded for the possibility that there may be too many individuals from other provinces. The red line represents $F_{st}$ threshold 0.05 for moderate genetic difference. Absolute difference between ancestry components 1 and 3 in different genomic loci (lower).

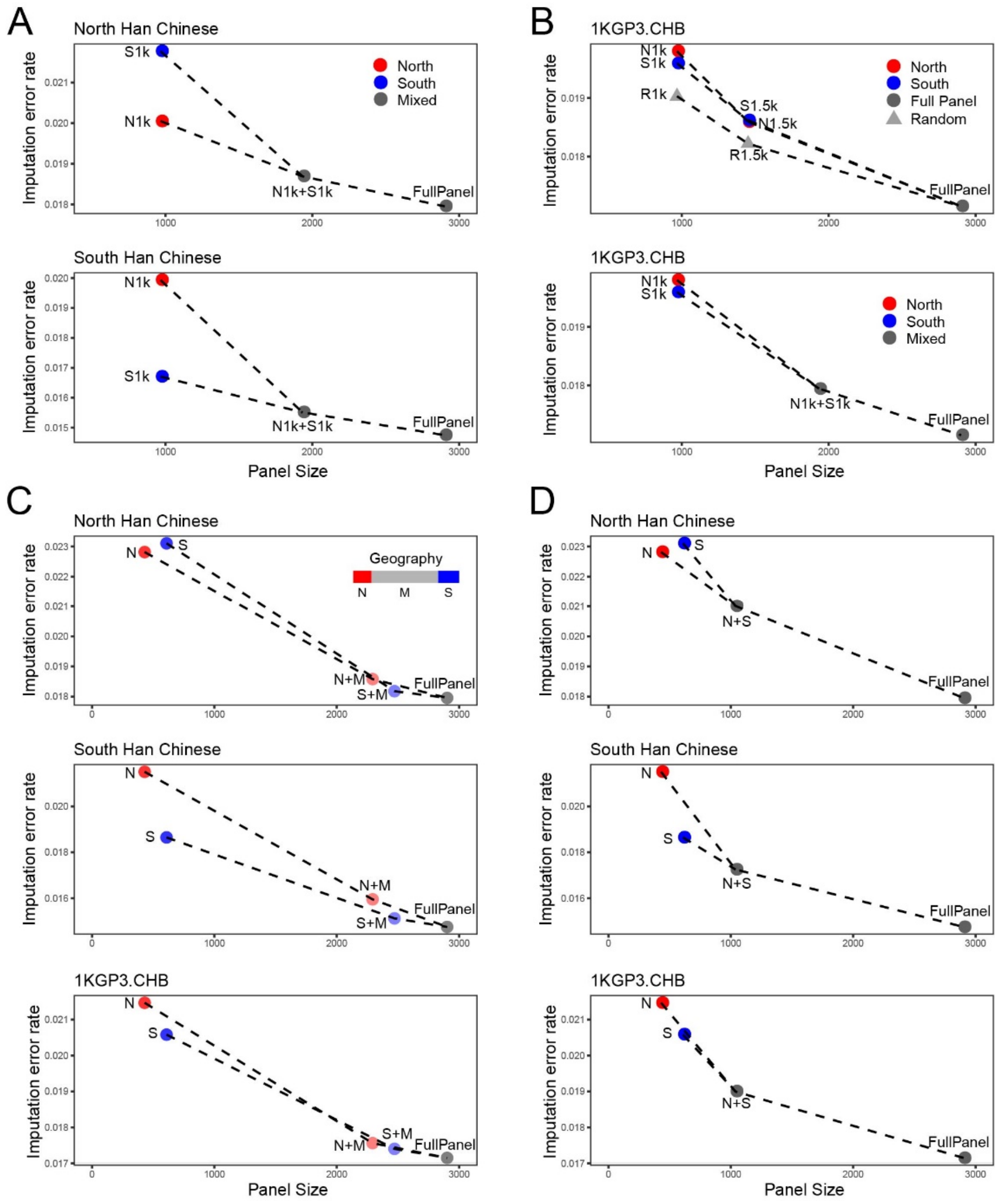


Figure S8. Imputation test for north and south samples using different panels.

(A) Imputation error rates for north (HGDP Han N. China, upper) and south (1KGP3 CHS, lower) Han Chinese using different subsets of NyuWa samples similar to Figure 4D. N1k+S1k means combining N1k and S1k samples as a mixed panel.

(B) Imputation error rates for 1KGP3 CHB samples. Tested panels are the same as Figure 4D (upper) and (A) (lower).

(C & D) Samples in NyuWa panel were classified as North (N) or South (S) using the classic geographical demarcation of Qinling Mountains-Huaihe River as in Figure S7C. Henan, Jiangsu, Anhui were excluded because the Huaihe River flows through these provinces. Shanghai was also excluded for the possibility that there may be too many individuals from other provinces. Samples in these provinces were grouped as Mixed (M). Imputation error rates for north (HGDP Han N. China, upper), south (1KGP3 CHS, middle) and 1KGP3 CHB (lower) Han Chinese were tested using different subsets and combinations of NyuWa samples. (C) Combining with mixed samples first. (D) Combining with unmatched samples first.

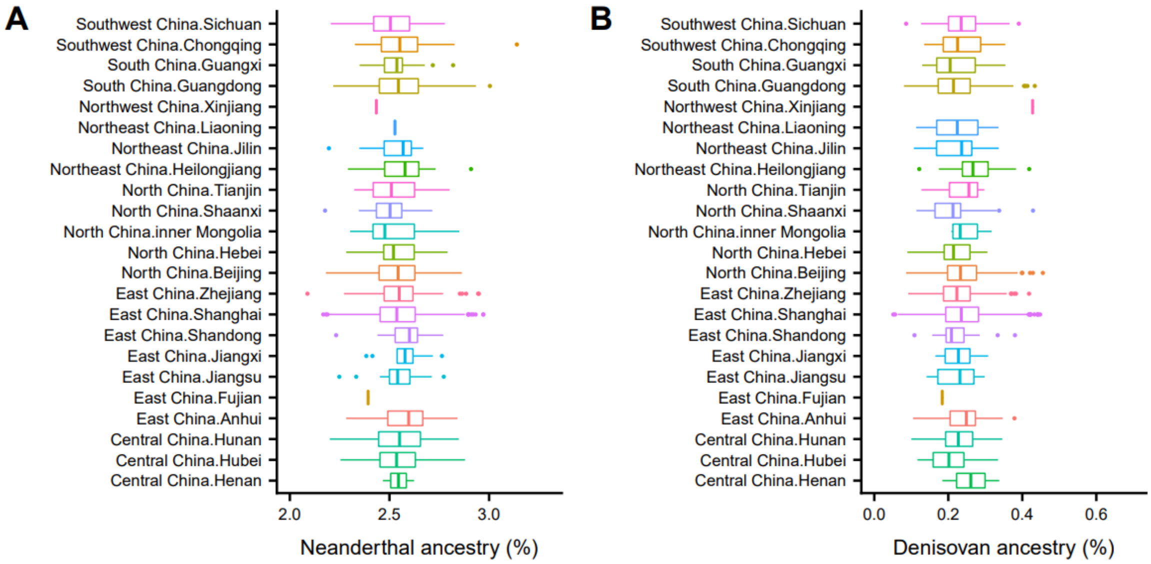


Figure S9. Introgression of Neanderthal (A) and Denisovan (B) ancestries in NyuWa samples.

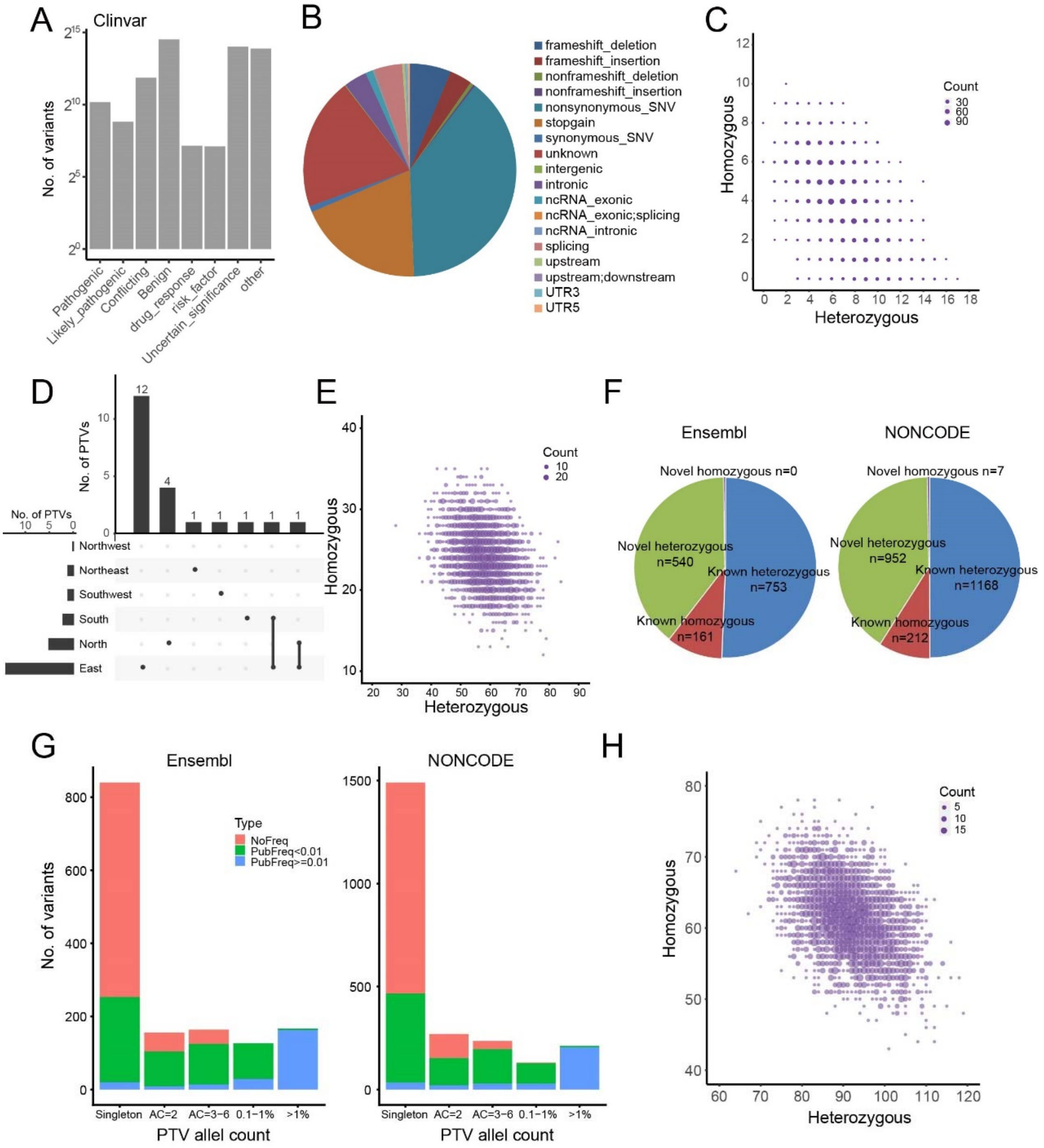


Figure S10. Clinical annotation and loss-of-function variants.

(A) Number of variants annotated as different categories by ClinVar.

(B) Distribution of ClinVar pathogenic variants in different types based on Ensembl annotation.

(C) Number of heterozygous and homozygous ClinVar pathogenic variants in NyuWa samples.

(D) Geographic regions of 25 novel homozygous PTVs in NyuWa samples.

(E) Number of heterozygous and homozygous PTVs in NyuWa samples.

(F) Number of lncRNA splicing variants by Ensembl and NONCODE annotations. Variants in both annotations are classified as Ensembl.

(G) Allele count/frequency distribution for lncRNA splicing variants by Ensembl and NONCODE annotations.

(H) Number of heterozygous and homozygous lncRNA splicing variants in NyuWa samples.
